# White matter connections of high-level visual areas predict cytoarchitecture better than category-selectivity in childhood, but not adulthood

**DOI:** 10.1101/2022.01.21.477131

**Authors:** Emily Kubota, Mareike Grotheer, Dawn Finzi, Vaidehi S. Natu, Jesse Gomez, Kalanit Grill-Spector

## Abstract

Ventral temporal cortex (VTC) consists of high-level visual regions that are arranged in consistent anatomical locations across individuals. This consistency has led to several hypotheses about the factors that constrain the functional organization of VTC. A prevailing theory is that white matter connections influence the organization of VTC, however, the nature of this constraint is unclear. Here, we test two hypotheses: (1) white matter tracts are specific for each category or (2) white matter tracts are specific to cytoarchitectonic areas of VTC. To test these hypotheses, we used diffusion magnetic resonance imaging (dMRI) to identify white matter tracts and functional magnetic resonance imaging (fMRI) to identify category-selective regions in VTC in children and adults. We find that in childhood, white matter connections are linked to cytoarchitecture rather than category-selectivity. In adulthood, however, white matter connections are linked to both cytoarchitecture and category-selectivity. These results suggest a rethinking of the view that category-selective regions in VTC have category-specific white matter connections early in development. Instead, these findings suggest that the neural hardware underlying the processing of categorical stimuli may be more domain-general than previously thought, particularly in childhood.

## Introduction

Ventral temporal cortex (VTC) consists of regions that respond more strongly to categories of stimuli such as faces (Kanwisher et al. 1997), places (Epstein and Kanwisher 1998), body parts (Peelen & Downing, 2005), and words (Cohen et al. 2003) than other stimuli (referred to as category-selective regions (Kanwisher 2010)). These regions fall in consistent anatomical locations across individuals (Grill-Spector and Weiner 2014; Weiner et al. 2014, 2018), which has led researchers to ask which factors constrain this functional organization. Classic theories suggest that function, cytoarchitecture, and connectivity all contribute to cortical organization (Zeki and Shipp 1988; Van Essen et al. 1992; Amunts and Zilles 2015). While past work has investigated the relationship between white matter and category-selectivity (Thomas et al. 2009; Saygin et al. 2011; Gschwind et al. 2012; Pyles et al. 2013; Yeatman et al. 2013; Bouhali et al. 2014; Gomez et al. 2015; Osher et al. 2016; Saygin et al. 2016; Grotheer et al. 2019; Li et al. 2020; Grotheer, Yeatman, et al. 2021) cytoarchitecture and category-selectivity (Weiner et al. 2017), and white matter and cytoarchitecture (Johansen-Berg et al. 2004; Klein et al. 2007), there has been limited research on the interplay between all three factors, which is the focus of the present study.

To investigate the relationship between white matter, cytoarchitecture, and category-selectivity, we use word-(mOTS-words, pOTS-words) and face-selective (pFus-faces and mFus-faces) functional regions of interest (ROIs) in VTC as a model system (**Figure 1A**). These regions are an excellent model system as mFus-faces and mOTS-words are located within the same cytoarchitectonic area, FG4, but are selective for different categories (faces and words, respectively). pFus-faces and pOTS-words are also located within the same cytoarchitectonic area, this time FG2, and are selective for words and faces, respectively (Weiner et al. 2017). That is, mOTS-words and pOTS-words are both selective for words, but are located within different cytoarchitectonic areas, and mFus-faces and pFus-faces are both selective for faces, but are located within different cytoarchitectonic areas. Notably, category-selectivity and cytoarchitecture are orthogonal in this case, allowing us to ask whether white matter connections align with category-selectivity, cytoarchitecture, or both.

**Figure 1.**
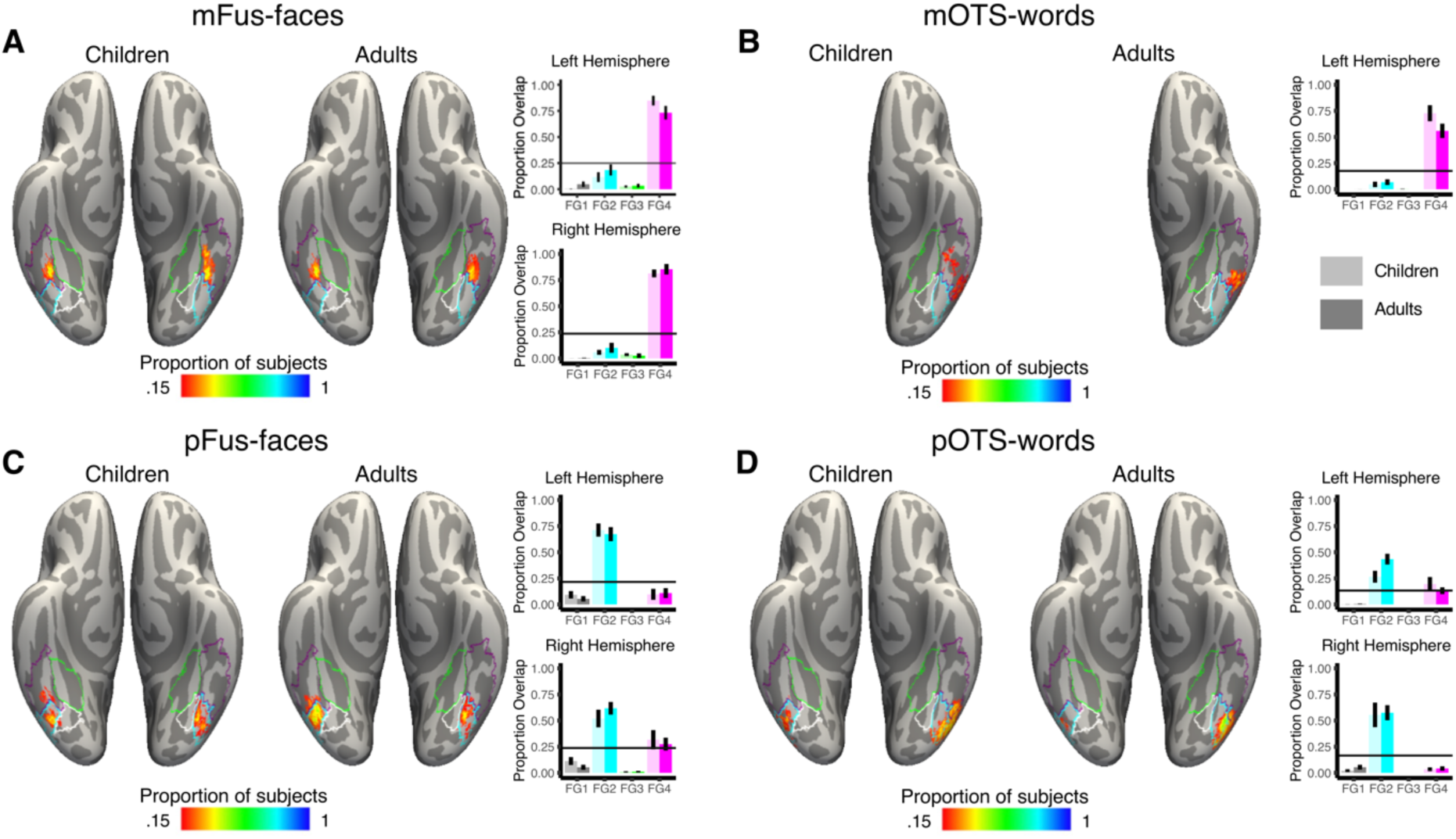
Children and adults have a similar functional-cytoarchitectonic relationship in ventral temporal cortex (VTC). Each brain panel shows the relationship between a probabilistic map of a functional region of interest (ROI) in children (left) and adults (right) vs. the maximal probability map (MPM) of the cytoarchitectonic areas of VTC on the FreeSurfer average ventral cortical surface (individual subject functional ROIs are shown in **Supplementary Figs 1-4**). *Vertex color*: proportion of subjects whose functional ROI includes that vertex (see colorbar). *Outlines:* MPM of cytoarchitectonic areas, FG1: white, FG2: cyan, FG3: green, FG4: purple. Bar graphs depict the proportion of overlap between each functional ROI and each of the 4 cytoarchitectonic areas. *Error bars:* standard error across subjects. *Lighter colors*: children, darker colors: adults. Upper: left hemisphere, lower: right hemisphere. (A) mFus-faces: mid fusiform face-selective region. (B) mOTS-words: mid occipitotemporal sulcus word-selective region. mOTS-words shown only for the left hemisphere as most subjects have this ROI only in the left hemisphere (C) pFus-faces: posterior fusiform face-selective region. (D) pOTS-words: posterior occipitotemporal sulcus word-selective region.

Previous studies provide support for the hypothesis that white matter connections of VTC functional regions are organized by category-selectivity. For example, studies examining the white matter fingerprint of VTC have shown that category-selective responses can be predicted by white matter connectivity to the rest of the brain (Saygin et al. 2011, 2016; Osher et al. 2016; Li et al. 2020). It has been hypothesized that different white matter connections of word- and face-selective regions may support the specific processing demands associated with each region’s preferred category. For example, word-selective regions may have more connections to language regions than face-selective regions to support the task of reading (Bouhali et al. 2014). This hypothesis aligns with the theory that white matter tracts might designate specific parts of cortex for processing different categories, perhaps from birth (Li et al. 2020). Supporting this view, researchers found that (i) congenitally blind subjects have a similar functional organization of VTC (van den Hurk et al. 2017; Murty et al. 2020) as well as similar functional connectivity to early visual cortex as sighted subjects (Butt et al. 2013; Striem-Amit et al. 2015), suggesting that there are innate anatomical constraints determining cortical organization (Mahon and Caramazza 2011), and (ii) the white matter fingerprint of VTC at age five can predict where word-selective cortex will emerge at age eight (Saygin et al. 2016). These studies suggest that white matter might be an innate or early developing constraint on category-selective processing. Moreover, these studies suggest that functional regions selective for the same category will have similar white matter connectivity to support category-specific processing.

Alternatively, classical investigations in human (Flechsig 1920; Zilles 2018) and nonhuman primates (Zeki and Shipp 1988; Van Essen et al. 1992) suggest that white matter connections may align with cytoarchitecture. For example, past work has shown that subregions of the frontal lobe with distinct cytoarchitecture also have distinct white matter connections (Johansen-Berg et al. 2004; Klein et al. 2007). Since the time of Brodmann, it has also been suggested that cytoarchitectonic areas might correspond to functional regions (Zilles 2018). However, recent work has demonstrated that this view is overly simplistic, as there is not always a one-to-one mapping between cytoarchitecture and functional regions (Glasser et al. 2016; Weiner, Barnett, et al. 2017; Rosenke et al. 2018). For example, a single cytoarchitectonic area in VTC, FG4, contains regions selective for three categories: faces, words, and body parts (Weiner, Barnett, et al. 2017). Therefore, these studies raise the possibility that functional regions located within the same cytoarchitectonic area might have similar white matter connectivity irrespective of category-selectivity.

In addition to different category-selective regions being located within the same cytoarchitectonic area, functional regions selective for the same category can be located in different cytoarchitectonic areas. For example, FG2 and FG4 each contain a word and face-selective region. These functional regions play different roles in the visual processing hierarchy (Grill-Spector and Weiner 2014; Grill-Spector et al. 2018) and therefore may have different white matter connections. Indeed, several studies have found that white matter connections of word-selective regions in VTC vary along the anterior-posterior axis of the ventral stream (Bouhali et al. 2014; Fan et al. 2014). Additionally, Lerma-Usabiaga et al. (2018) found that the word-selective region in FG2 has different white matter connections and functional properties compared to the word-selective region in FG4. While these studies suggest that white matter connectivity varies along the anterior-posterior axis of the ventral stream for word-selective regions, it is unclear if the same is true for face-selective regions. Past work demonstrated that there are local white matter connections between regions constituting the face-processing network (Gschwind et al. 2012; Pyles et al. 2013), suggesting that these regions may have distinct long range connections. Together, these data raise the possibility that functional regions selective for the same category may have distinct cytoarchitecture and white matter connections to support distinct processing demands across the visual processing hierarchy.

In addition to the debate on whether the white matter connections of VTC functional ROIs are category-specific or cytoarchitectonic-specific, an important question is whether the white matter connections of VTC are fixed across development or if they change with the functional development of VTC during childhood. Indeed, while some functional regions, like face-selective regions may be present in infancy (Kosakowski et al. 2021), other regions, such as word-selective regions, emerge later, only when children learn to read (Brem et al. 2010; Dehaene-Lambertz et al. 2018; Kubota et al. 2019). Notably, both word- and face-selective regions continue to develop well into adolescence (Turkeltaub et al. 2003; Scherf et al. 2007; Golarai et al. 2010; Ben-Shachar et al. 2011; Cantlon et al. 2011; Dehaene-Lambertz et al. 2018; Kubota et al. 2019; Nordt et al. 2021). We reasoned that if white matter connections lay an intrinsic blueprint for later development of category-selectivity (Bouhali et al. 2014; Saygin et al. 2016; Li et al. 2020), then the white matter connections of VTC functional ROIs will be consistent across development. This hypothesis is supported by evidence that the connectivity fingerprint of VTC category-selective regions seems to be stable during the first year of schooling (Moulton et al. 2019) and that white matter connections in 5-year-olds predict VTC word-selective activations in 8-year-olds (Saygin et al. 2016). Alternatively, as category-selective regions continue to develop from childhood into adolescence, there may be a dynamic interplay between the development of functional regions and their white matter connectivity, especially considering the protracted development of word and face-selective regions. This hypothesis predicts that white matter connections of face- and word-selective regions will change from childhood to adulthood.

To determine the nature of VTC white matter connections and their development, we collected functional magnetic resonance imaging (fMRI) and diffusion magnetic resonance imaging (dMRI) in 27 children (ages 5.76 -12.39 years, mean±sd: 9.74±2.11 years, 15 female, 12 male) and 28 adults (22.08 – 28.58 years, 24.09±1.59 years, 11 female, 17 male). We used a fMRI localizer experiment (Stigliani et al. 2015) to define functional ROIs in each individual’s brain and dMRI data to generate a whole brain white matter connectome for each individual. We intersected each subject’s whole brain connectome with their functional ROIs to identify the functionally defined white matter associated with each functional ROI (Grotheer et al. 2019; Finzi et al. 2021; Grotheer, Yeatman, et al. 2021). We then tested if the functionally defined white matter of face (pFus- and mFus-faces) and word-selective regions (pOTS- and mOTS-words) is linked to category, cytoarchitecture, or both, and if they change from childhood to adulthood.

## Materials and Methods

### Subjects

The study was approved by the Institutional Review Board of Stanford University. Prior to the start of the study, all subjects gave written consent. 29 children and 28 adults participated in the experiment. One child was excluded for an incomplete dataset, and one child was excluded due to a cyst on the anterior temporal lobe. Thus, we report data from N = 27 children (ages 5.76 - 12.39 years, mean = 9.74, standard deviation = 2.11; 15 female, 12 male), and N=28 adults (ages 22.08 – 28.58 years, mean=24.09, standard deviation = 1.59; 11 female, 17 male). The group sizes and age range are consistent with other studies from our lab finding developmental effects in face and word-selective regions (Gomez et al. 2018; Natu et al. 2019; Nordt et al. 2021). The racial demographics of the sample were as follows: 56% White, 15% Asian, 9% mixed-race, 7% Hispanic, 5% Black, 4% other, 4% missing data.

## MRI

MRI data were collected at the Center for Cognitive and Neurobiological Imaging at Stanford University, using a GE 3 Tesla Signa Scanner with a 32-channel head coil.

### Anatomical MRI

We acquired quantitative MRI measurements using protocols from Mezer et al. (2013) to generate a 1mm isotropic T1-weighted anatomical image of each subject. Anatomical images were used for gray matter/white matter segmentation, cortical surface reconstruction, and visualization of data on the cortical surface.

### Diffusion MRI (dMRI) acquisition

Diffusion-weighted dMRI was acquired using a dual-spin echo sequence in 96 different directions, 8 non-diffusion-weighted images were collected, 60 slices provided full head coverage. Voxel size=2 mm isotropic, repetition time (TR)=8000ms, echo time (TE)=93.6ms, field of view (FOV)=220mm, flip angle=90°, b0=2000 s/mm^2^.

### Functional MRI acquisition

We acquired simultaneous multi-slice, one-shot T2* sensitive gradient echo EPI sequences with a multiplexing factor of 3 and near whole brain coverage (48 slices), FOV=192mm, TR=1s, TE=30ms, and flip angle=76°. Voxel size=2.4 mm isotropic.

#### Functional localizer task

Subjects took part in a 10-category functional localizer experiment (Stigliani et al. 2015; Nordt et al. 2021). The stimuli included faces (children, adults), characters (pseudowords, numbers), bodies (headless bodies, limbs), places (houses, corridors), and objects (cars, string instruments). Stimuli were presented in 4 second blocks with 2 Hz stimulus presentation rate. Subjects were instructed to fixate throughout the experiment and complete an oddball task, which required pressing a button whenever a phase-scrambled image appeared. Each run was 5 minutes and 18 seconds long and subjects completed 3 runs of the experiment.

## Data Analyses

All analyses were done in the individual brain space of each subject.

### Diffusion MRI processing

dMRI data were preprocessed using MRtrix3 (Tournier et al., 2019; https://github.com/MRtrix3/mrtrix3) and mrDiffusion (http://github.com/vistalab) as in past work (Grotheer et al., 2019; https://github.com/VPNL/fat). Data were denoised using principal component analysis, Rician based denoising, and Gibbs ringing corrections. Eddy currents and motion were corrected for using FSL (https://fsl.fmrib.ox.ac.uk/), and bias correction was performed using ANTs (Tustison et al. 2010). dMRI data were aligned to the b0 images and then aligned to the high-resolution T1-weighted image of the same brain using a six-parameter rigid-body alignment. Voxel-wise fiber orientation distributions (FOD) were calculated using constrained spherical deconvolution (CSD) (Tournier et al. 2007).

### dMRI exclusion criterion

The exclusion criterion for the dMRI data was data with more than 5% outliers (intra-volume slices affected by signal dropout) identified with FSL’s eddy tool (https://fsl.fmrib.ox.ac.uk/fsl/fslwiki/eddy). No subjects had more than 5% outliers, thus, all subjects were included under this criterion.

### Generating white matter connectomes

We used MRtrix3 to generate five candidate connectomes with the following maximum angles (2.25°, 4.5°, 9°, 11.25°, and 13.5°). The goal of using multiple candidate connectomes was to avoid biasing the results towards either long range or local connections (Takemura et al. 2016). For each candidate connectome we used probabilistic tracking with the IFOD1 algorithm, a step size of 0.2 mm, a minimum length of 4 mm, a maximum length of 200 mm, and a fiber orientation distribution (FOD) amplitude stopping criterion of 0.1. Each candidate connectome consisted of 500,000 streamlines. We used anatomically constrained tractography (ACT; Smith et al., 2012) to identify the gray-matter-white-matter interface directly adjacent to the functional ROIs. Streamlines were randomly seeded on this gray-matter-white-matter interface. This procedure is critical for accurately identifying the connections that reach functional ROIs, which are located in the gray matter (Grotheer, Kubota, et al. 2021). Random seeding with a constant number of streamlines results in a different seed density across subjects as brain size differs across individuals. Finally, we concatenated the candidate connectomes into a single connectome containing 2,500,000 streamlines. We refer to this as the white matter connectome of each subject.

To identify the main white matter bundles (fascicles) of the brain we used Automated Fiber Quantification (Yeatman, Dougherty, Myall, et al. 2012) (AFQ; https://github.com/yeatmanlab/AFQ). The output of AFQ is 11 fascicles in each hemisphere and 2 cross-hemispheric fascicles.

### fMRI processing

Functional data was aligned to the anatomical brain volume of each individual subject. Motion correction was performed both within and across runs. No spatial smoothing or slice-timing correction was performed. To estimate the response to each category a general linear model (GLM) was fit to each voxel by convolving the stimulus presentation with the hemodynamic response function.

### fMRI exclusion criterion

Criteria for exclusion of data were within-run motion >2 voxels or between-run motion >3 voxels. All subjects were included under this threshold.

### Functional region of interest definition

To identify a functional ROI that showed significant selectivity to exemplars of a particular category, we used a contrast comparing the response to the category of interest compared to the other categories and included voxels that passed the statistical threshold (t > 3, voxel level uncorrected). For word-selective regions mOTS-words and pOTS-words, we contrasted words > all other categories (excluding numbers). For faces, we contrasted adult and child faces > all others. We defined the functional ROIs on the inflated cortical surface of each subject to ensure the correct anatomical location of each functional ROI.

### Determining the relationship between functional ROIs and cytoarchitectonic areas

To determine the relationship between functional ROIs and cytoarchitectonic areas we registered each individual’s functional ROIs to the fsaverage cortical surface and measured the overlap between each functional ROI and a probabilistic estimate of each of the cytoarchitectonic areas of ventral temporal cortex (FG1-FG4) on the fsaverage cortical surface (Rosenke et al. 2018). The Rosenke 2018 Atlas contains the maximal probabilistic maps of 8 cytoarchitectonic areas of the human ventral visual stream (hOc1-hOc4; FG1-FG4) generated from observer-independent definitions of cytoarchitectonic areas by the Amunts and Zilles labs (Amunts et al. 2000; Rottschy et al. 2007; Caspers et al. 2013; Lorenz et al. 2017). To quantify the overlap between a functional ROI and a cytoarchitectonic area, we calculated the number of vertices shared between a given functional ROI and a cytoarchitectonic area divided by the total number of vertices belonging to that functional ROI (and then multiplied by 100 to get a percentage). To calculate the chance level overlap for each functional ROI, we calculated the overlap between each functional ROI in each subject and a randomly selected cytoarchitectonic area with replacement 400 times and then calculated the mean overlap (as in Weiner et al., 2017).

### Connectivity profiles of functionally defined white matter

To identify the white matter associated with each functional region we intersected each functional ROI (mOTS-words, mFus-faces, pOTS-words, pFus-faces; defined from the fMRI data) with the white matter connectome, the product of which we refer to as *functionally defined white matter*. All analyses were done in the native brain space of each subject. For each functional ROI we defined a connectivity profile in two ways: (1) its fascicle connectivity profile, and (2) its endpoint connectivity profile.

#### (1) Fascicle connectivity profile

To identify the fascicles connected to each functional ROI, we intersected the subject’s fascicles classified by AFQ with the gray matter/white matter interface directly adjacent to that functional ROI (see Grotheer et al. 2019; Finzi et al. 2021; Grotheer, Kubota, et al. 2021). This process gave us the functionally defined white matter fascicles for each functional ROI. Previous work showed that out of those fascicles classified by AFQ, only 5 fascicles connect to VTC: the inferior longitudinal fasciculus (ILF), inferior frontal occipital fasciculus (IFOF), arcuate fasciculus (AF), posterior arcuate fasciculus (pAF), and vertical occipital fasciculus (VOF) (Ben-Shachar et al. 2007; Thomas et al. 2009; Wandell and Yeatman 2013; Yeatman et al. 2013; Tavor et al. 2014; Gomez et al. 2015; Song et al. 2015; Weiner et al. 2016; Kay and Yeatman 2017; Lerma-Usabiaga et al. 2018; Broce et al. 2019; Grotheer et al. 2019; Grotheer, Yeatman, et al. 2021). To determine the fascicle connectivity profile, we calculated the distribution of its connections across these five fascicles for each functional ROI in each subject. That is, the percentage of streamlines belonging to each fascicle out of the total number of streamlines connecting to the functional ROI.

#### (2) Endpoint connectivity profile

To identify the endpoints associated with each functional ROI, we intersected the whole brain connectome of each subject with the gray matter/white matter interface directly adjacent to the functional ROI. This approach identifies all white matter tracts that intersect with the functional ROI and not only those which belong to the main fascicles classified by AFQ. We projected the endpoints of the functionally defined white matter to the cortical surface using tract density imaging (TDI) with mrTrix3 (Tournier et al., 2019), and calculated the distribution of endpoints of these tracts across all cortical surface vertices. We transformed the TDI output into a distribution of endpoints by dividing the endpoint map by the total number of endpoints. This results in an endpoint density map that sums to one for each functional ROI and subject. To quantify the endpoint density map in an interpretable manner that is related to a common whole brain atlas, we measured the average endpoint density in each of the 180 (per hemisphere) Glasser Atlas ROIs (Glasser et al. 2016). Thus, we projected the Glasser Atlas from the fsaverage cortical surface into each subject’s native cortical brain surface using cortex-based alignment in FreeSurfer version 6.0, and then calculated in each subject’s cortical surface the average endpoint density in each of the Glasser ROIs. We also calculated the average endpoint connectivity density maps across subjects by projecting each individual’s subject map to the fsaverage cortical surface using cortex-based alignment and averaging across subjects.

### Evaluating similarity between connectivity profiles

We used a within-subject correlation metric to calculate the similarity between connectivity profiles of pairs of functional ROIs. That is, for each pair of ROIs we calculated the correlation between connectivity profiles in each subject with both ROIs defined. To report the effect size, all correlation values in the figures show the Pearson correlation values. To ensure normality of the data when calculating statistical significance, all statistics are conducted on Fisher transformed correlation values (Fisher 1915).

### Distance control ROI definition

To control for differing distances between ROIs within and across cytoarchitectonic areas, we performed two control analyses: (1) we included distance as factor in statistical analyses (see Statistics section below), and (2) we examined if equidistant ROIs within the same cytoarchitectonic area or across cytoarchitectonic area would have equally similar connectivity profiles. To that end, we drew equidistant control ROIs on the fsaverage surface. The goal was to generate three ROIs with the center ROI equidistant from the other two ROIs, one of which was located within the same cytoarchitectonic area and the other of which was located in a different cytoarchitectonic area. We chose these control ROIs along a posterior to anterior axis as this axis is consistent with the spatial arrangement of FG2 and FG4 and is the elongated anatomical axis. Control ROIs were defined using anatomy alone, using an iterative process as the cytoarchitectonic ROIs are on the fsaverage cortical surface and we calculated the Euclidean distance between the ROIs in individual subject volumetric space, since the white matter meets the gray matter in volumetric space. First, we drew ROIs on the fsaverage cortical surface with the following constraints: 1) two ROIs located in FG4 and one ROI located in FG2 and 2) as close to the center of the cytoarchitectonic ROIs as possible as the cytoarchitectonic ROIs are probabilistic maps and may have more variability in the edges of individual subjects. Next, we transformed the ROIs to volumetric space and calculated the Euclidean distance between the center ROI and the ROI located within the same cytoarchitectonic area, and the Euclidean distance between the center ROI and the ROI located within a different cytoarchitectonic area. If the distance between the ROIs was still significantly different then we re-drew the ROIs on the fsaverage space and repeated the process until the ROIs were equidistant.

### Predicting cortical parcellations from connectivity profiles

To test whether connectivity profiles predict the cytoarchitectonic area, the category-selectivity of a functional ROI, or both, we used a winner-takes-all maximum correlation classifier using a leave-one-out cross-validation procedure. All three classification types were performed for both the fascicle connectivity profiles and the endpoint connectivity profiles.

a. *Cytoarchitecture classification (2-way classification): Training*: We calculated the average connectivity profile of functional ROIs in each cytoarchitectonic area (FG2 and FG4), excluding the held-out subject’s connectivity profiles. *Testing*: We classified the cytoarchitectonic area of each of the unlabeled, held-out connectivity profiles based on the highest correlation with the connectivity profiles of the training set. That is, the held-out connectivity profile was classified as belonging to the cytoarchitectonic area with which it was most highly correlated. This classification was then compared to the true cytoarchitectonic area to determine the accuracy of the classifier. We repeated this procedure across all n-folds for the connectivity profile of each functional ROI.
b. *Category classification (2-way classification): Training:* We calculated the average connectivity profile of functional ROIs with the same category-selectivity (words and faces), excluding the held-out subject’s connectivity profiles. *Testing*: We classified the cytoarchitectonic area of each of the unlabeled, held-out connectivity profiles based on the highest correlation with the training connectivity profiles. This classification was then compared to the true category-selectivity to determine the accuracy of the classifier. We repeated this procedure across all n-folds for the connectivity profile of each functional ROI.
c. *Functional ROI classification (4-way classification): Training:* We calculated the average connectivity profile for each functional ROI (mFus-faces/mOTS-words/pFus-faces/pOTS-words) excluding the held-out subject’s connectivity profiles. *Testing:* We classified the connectivity profile of the held-out functional ROI based on the highest correlation with the training connectivity profiles. This classification was then compared to the true functional ROI label to determine the accuracy of the classifier. We then repeated the procedure for each connectivity profile of each functional ROI across all n-folds.

### Controls for development differences

As we found developmental differences, we further examined whether between age-group differences in motion, functional ROI size, or functional ROI location could drive the developmental differences in connectivity profiles across children and adults.

### Motion control

Even though all subjects met our inclusion criterion, both rotation and translation were significantly different between children and adults (rotation: *t*(53) = -2.63, *p* = 0.01, translation: *t*(53) = -2.24, *p* = 0.03). To control for this difference, we repeated the analyses comparing across age-groups (connectivity profile similarity analysis and classification analysis) using the subset of subjects that were matched across ages for motion. This was achieved by excluding 5 subjects that had either the most rotation (n=4) or translation (n=2, one overlapping with rotation). The excluded subjects were: one six-year-old, one seven-year-old, two nine-year-olds, and one eleven-year-old. After removing these five subjects the mean age of the children was 9.87± 2.16 years and there were no longer significant differences in rotation (*t*(48) = -1.67, *p* = 0.10) and translation (*t*(48) = -0.98, *p* = 0.33) across children and adults.

### Constant size ROI control

For the main analyses, we use functional ROIs defined from contrasts in each subject’s brain as outlined above because it is the most accurate way to define functional ROIs for each subject. However, if there are developmental changes in ROIs sizes, they may affect connectivity profiles across age groups. Therefore, as a control, we repeated both fascicle and endpoint connectivity analyses using a constant sized, 3mm disk ROI centered at the center of each functional ROI on each subject’s native brain.

### Smaller age range control

As there is a large age range in our child subjects (ages 5.76 -12.39 years, mean = 9.74, standard deviation = 2.11), we repeated the analyses with a smaller age range from 9-11 years old. This group included 18 child subjects (ages 9.18 -11.97 years, mean = 10.66, standard deviation = 0.92).

### VTC exclusion control

For the main analyses of white matter endpoints, we examined all endpoints across the entire brain as this reflects the entirety of the connectivity endpoint maps. However, as some of these endpoints are in VTC, they may be particularly affected by between age group differences in the location and spatial extent of functional ROIs within VTC. To further test that development of endpoint connectivity is not just due to developmental differences restricted to VTC, we repeated connectivity endpoints analyses excluding 11 Glasser ROIs that overlap with VTC (see Nordt et al., 2019): PHA1, PHA3, TE2p, PH, PIT, FFC, V8, VVC, VMV2, VMV3, TF. Note that these are the Glasser ROIs naming conventions rather than the naming conventions in visual neuroscience (Brewer et al. 2005; Wandell and Winawer 2011; Wang et al. 2015; Rosenke et al. 2021).

### Statistics

In general, statistical analyses were computed using the lmerTest (Kuznetsova et al. 2017) package in R. Linear mixed-effects models (LMMs) were used because our data include missing data points (not all subjects have all functional ROIs). The significance of LMM model terms was evaluated using repeated-measures analyses-of-variance (LMM ANOVAs; type III) with Satterthwaite’s method of correction for degrees of freedom (Luke 2017).

### Testing for significant differences in similarity among connectivity profiles

We measured the pairwise correlation between connectivity profiles to quantify their similarity. To test whether connectivity profiles were more correlated for functional ROIs located within the same cytoarchitectonic area or selective for the same category and if this differed by age group, we used the following model:

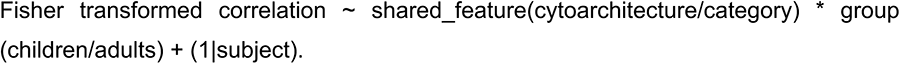

To test if distance is a factor, we also considered an additional model:

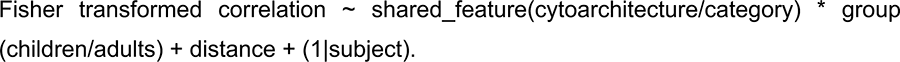

Log-likelihood testing revealed that distance is a significant independent factor (p=1.25*10^−5^ for fascicle connectivity and p=5.83*10^−9^ for endpoint connectivity).

### Testing if classification accuracy is significantly above chance

To test whether classification accuracy was above chance, we performed a one-sample *t*-test testing whether classification was significantly different than the chance level (50% for cytoarchitecture/category classification; 25% for functional ROI classification).

### Testing for significant differences in classification accuracy

To test whether classification accuracy differed by classification type (cytoarchitecture/category) and age group (children/adults) we performed a binomial logistic regression with the following model: Classification accuracy ∼ classification feature (cytoarchitecture/category) * group (children/adults). We then assessed the odds ratio and its confidence interval (CI). Odd ratios are significant if 1 is outside the 95% CI.

### Testing whether connectivity profiles are more correlated for ROIs located in the same cytoarchitectonic area, when controlling for position along the anterior-posterior axis

To test whether connectivity profiles were more similar when they were located within the same cytoarchitectonic area, even when location along the anterior-posterior axis was held constant, and whether this differed across age groups we used the following model: Fisher transformed correlation ∼ cytoarchitecture(same/different) * group (children/adults) + (1|subject).

## Results

### Testing the relationship between functional ROIs and cytoarchitecture in children and adults

Before testing if white matter connectivity is linked to cytoarchitecture or category-selectivity, we tested whether there is a consistent relationship between functional ROIs and cytoarchitectonic areas in children and adults and if this is similar across age groups. To assess this relationship, we followed the procedure of Weiner et al. (2017) in which we quantified the overlap between each functional ROI and each probabilistic cytoarchitectonic area (FG1, FG2, FG3, FG4) from the Rosenke Atlas (Rosenke et al. 2018) in each subject and compared across ROIs and age groups.

We were able to identify functional ROIs in most subjects (Left hemisphere: children: mFus-faces: n = 22, mOTS-words: n = 20, pFus-faces, n = 20, pOTS-words: n= 17; adults: mFus-faces: n = 26, mOTS-words: n = 25, pFus-faces, n = 20, pOTS-words: n = 26. Right hemisphere: children: mFus-faces: n = 23, pFus-faces, n = 18, pOTS-words: n= 12; adults: mFus-faces: n = 21, pFus-faces, n = 23, pOTS-words: n = 24).

**Fig 1** shows the probabilistic maps of functional ROIs with respect to the cytoarchitecture in child and adult subjects, demonstrating three main findings. First, in both children and adults both the functional ROIs and cytoarchitectonic areas align to previously established anatomical landmarks. For example, mFus-faces is located at the anterior end of the mid-fusiform sulcus (Weiner et al. 2014). Second, we replicate the findings of Weiner et al. (2017) in a new set of adult subjects. That is, we find that mFus-faces and mOTS-words are located in FG4 with above chance frequency, whereas pFus-faces and pOTS-words are located in FG2 above chance. Third, we find that this functional-cytoarchitectonic relationship is also observed in children. To quantify the significance of this relationship, for each subject and functional ROI, we quantified the proportion overlap between that ROI and each of the four cytoarchitectonic areas (bar graphs in **Fig 1**). Then we used a linear model with factors of cytoarchitectonic area, age group, and hemisphere (with the exception of mOTS-words which is only in the left hemisphere) to test for significant differences.

In both children and adults, mFus-faces and mOTS-words were mostly within FG4 (**Fig 1A,B**). Indeed, there was a main effect of cytoarchitectonic area (mFus-faces: F(3,352) = 436.06, p < 2×10^−16^; mOTS-words: F(3,172) = 127.40, p < 2×10^−16^). All other main effects and interactions were not significant (ps > .06, Fs < 2.58) suggesting that functional-cytoarchitectonic coupling was not different across age groups.

For pFus-faces and pOTS-words, there was also a main effect of cytoarchitectonic area reflecting greater overlap with FG2 compared to the other cytoarchitectonic ROIs (pFus-faces: *F*(3,308) = 129.71, *p* < 2.2×10^−16^; pOTS-words: *F*(3,300) = 110.04, *p* < 2.2 ×10^−16^, **Fig 1C,D**). There was also a hemisphere by cytoarchitectonic area interaction (pFus-faces: *F*(3,308) = 6.60, *p* = 0.0003); pOTS-words: *F*(3,300) = 10.46, *p* = 1.46×10^−6^), where there was greater correspondence between the functional ROIs and cytoarchitectonic area FG2 in the right hemisphere. All other main effects and interactions were not significant, reflecting no significant differences between age groups (*p*s > 0.13, *F*s < 2.23). The observed hemispheric difference likely reflects hemispheric differences in the probabilistic cytoarchitectonic area definitions, where FG2 extends more laterally in the right hemisphere compared to the left hemisphere. As the cytoarchitectonic areas were identified using data from 10 postmortem brains (Caspers et al. 2013), it is unknown whether this observed hemispheric asymmetry would hold in a larger sample.

As these analyses validate that the functional-cytoarchitectonic structure of VTC is similar across children and adults, we next examined the functionally defined white matter for face- and word-selective VTC ROIs, using two complementary approaches. In the first approach, we identified the fascicles (Catani et al. 2002; Yeatman, Dougherty, Myall, et al. 2012) that connect to each functional ROI, and calculated the distribution of connections across the fascicles. We refer to this quantification as the *fascicle connectivity profile* of each functional region. The fascicle connectivity profile provides an interpretable quantification of the white matter connections of each functional ROI because the anatomical layout of the major fascicles is well-established (Catani et al. 2002; Wakana et al. 2004; Yeatman, Dougherty, Myall, et al. 2012). In the second approach, which we refer to as the *endpoint connectivity profile*, we determined the distribution of endpoints of the white matter connections of each functional ROI across the entire cortex.

### Fascicle connectivity profiles of VTC functional ROIs are organized by cytoarchitecture in children & adults

We first examined the fascicle connectivity profiles of VTC face- and word-selective regions in each individual’s brain. There are five fascicles that connect to visual regions in VTC (Ben-Shachar et al. 2007; Thomas et al. 2009; Wandell and Yeatman 2013; Yeatman et al. 2013; Tavor et al. 2014; Gomez et al. 2015; Song et al. 2015; Weiner et al. 2016; Kay and Yeatman 2017; Lerma-Usabiaga et al. 2018; Broce et al. 2019; Grotheer et al. 2019; Grotheer, Yeatman, et al. 2021) (**Fig 2A**-top): the inferior frontal occipital fasciculus (IFOF) (Catani et al. 2002), which connects the occipital lobe and frontal lobe, the inferior longitudinal fasciculus (ILF) (Catani et al. 2002), which connects the occipital lobe and ventral temporal lobe, the arcuate fasciculus (AF) (Catani et al. 2002), which connects the ventral temporal lobe and the frontal lobe, the posterior arcuate fasciculus (pAF) (Weiner, Yeatman, et al. 2017), which connects the ventral temporal lobe and the parietal lobe, and the vertical occipital fasciculus (VOF) (Yeatman, Weiner, et al. 2014; Takemura et al. 2016; Weiner, Yeatman, et al. 2017), which connects the ventral and dorsal occipital lobe. Therefore, our fascicle connectivity analysis will focus on these five fascicles. Since word-selective regions are left lateralized (Petersen et al. 1988), left hemisphere data are provided in the main figures and right hemisphere data are presented in the supplemental figures.

**Figure 2.**
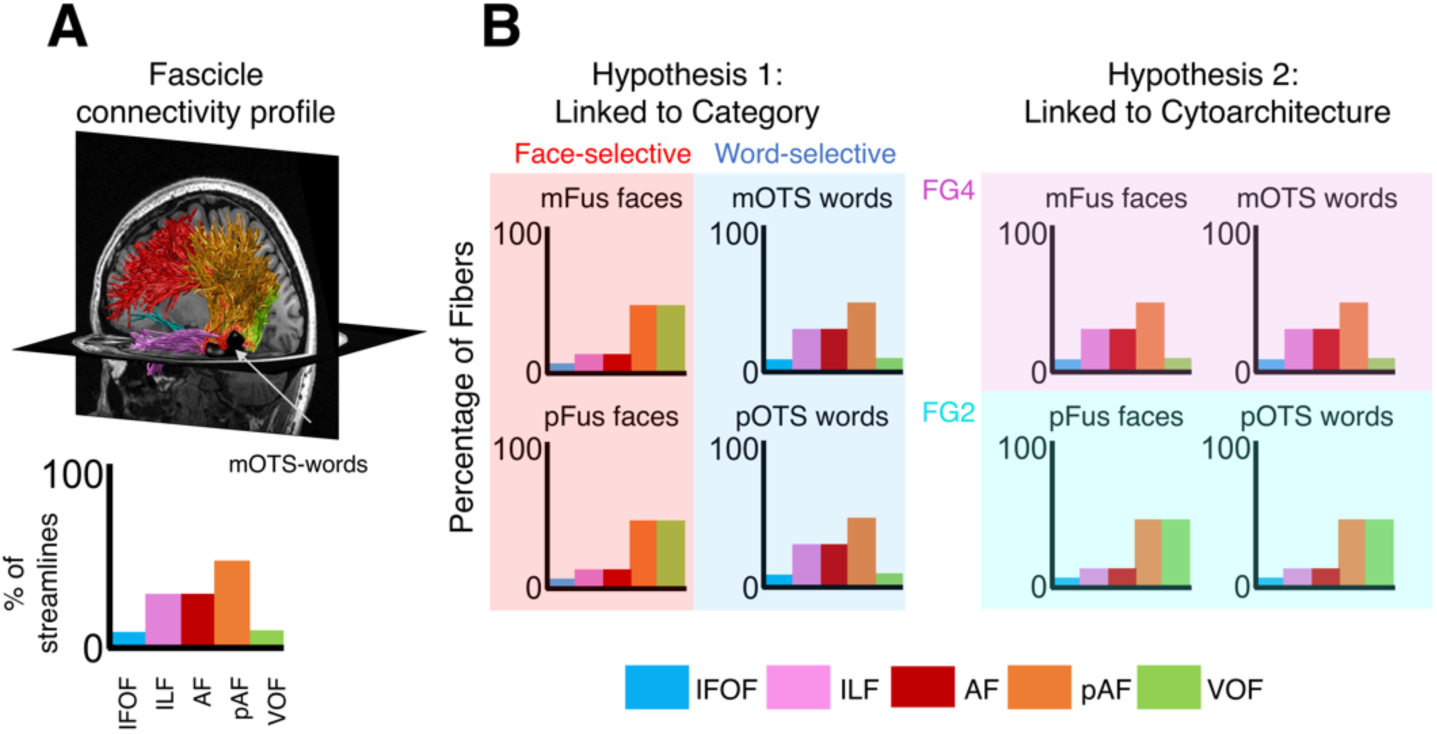
Design and hypotheses. **(A)** *Top*: example fascicle connectivity profile in a single participant for a word-selective region, mOTS-words (black). *Bottom*: Quantification of the connectivity profile shown above: the percentage of fibers connecting to these five fascicles. (B) Schematic of two opposing hypotheses relating the organization of white matter tracts to functional regions. Each panel shows the predicted distributions of connections across white matter fascicles for a given functional ROI. *Bar Colors:* fascicles. *Acronyms:* pFus: posterior fusiform; mFus: mid fusiform; pOTS: posterior occipito-temporal sulcus. mOTS: mid occipito-temporal sulcus. IFOF: inferior fronto-occipital fasciculus; ILF: inferior longitudinal fasciculus; AF: arcuate fasciculus; pAF: posterior arcuate fasciculus; VOF; vertical occipital fasciculus.

We reasoned that if the white matter connections of ventral visual regions are linked to category-selectivity, then we would expect that (i) mFus-faces and pFus-faces would have similar white matter connections to support face recognition, (ii) mOTS-words and pOTS-words would have similar white matter connections to support word recognition, and (iii) white matter connections of word-selective regions would be different from those of face-selective regions (**Fig 2B, Hypothesis 1**). However, if the white matter connections of VTC are organized by cytoarchitecture, then we would expect (i) that mFus-faces and mOTS-words would have similar white matter connections as they reside in the same cytoarchitectonic area (FG4), (ii) that pFus-faces and pOTS-words would have similar white matter connections to each other (as they reside in FG2), and (iii) that the connectivity profile of regions in FG2 (pFus-faces and pOTS-worlds) will be different from those in FG4 (mFus-faces and mOTS-words) irrespective of category-selectivity (**Fig 2B, Hypothesis 2**).

In both children and adults, ventral face- and word-selective regions showed a nonuniform fascicle connectivity profile (**Fig 3**). Qualitatively, pFus-faces and pOTS-words showed prominent connections to the VOF, and mFus-faces and mOTS-words showed prominent connections to the pAF (example child in **Fig 3A;** all children in **Supplemental Figs 5-8**, all adults in **Supplemental Figs 9-12**).

**Figure 3.**
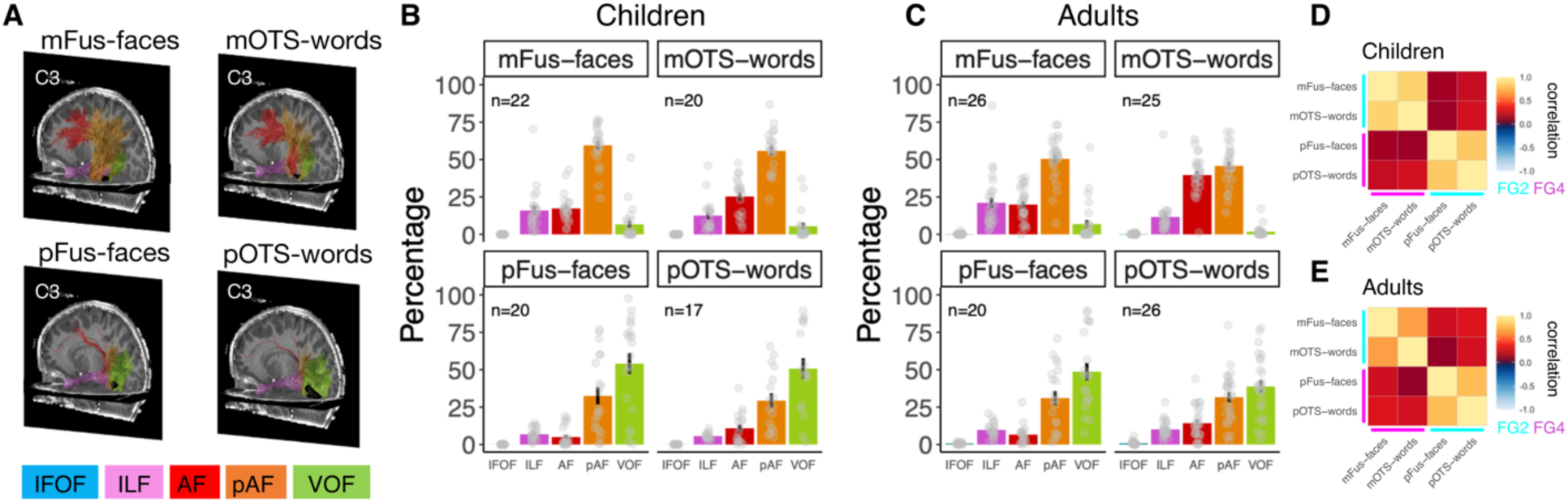
Fascicle connectivity profiles of VTC face- and word-selective regions. **(A)** Fascicle connectivity profiles of face- and word-selective functional ROIs in an example child (C3, 6-years-old). All subjects in **Supplemental Figs 5-12**. *Acronyms: mFus-faces:* mid fusiform face-selective region. *pFus-faces:* posterior fusiform face-selective region. *mOTS-words:* mid occipitotemporal sulcus word-selective region *IFOF:* inferior fronto-occipital fasciculus; *ILF:* inferior longitudinal fasciculus; *AF:* arcuate fasciculus; *pAF:* posterior arcuate fasciculus; *VOF:* vertical occipital fasciculus. (B,C) Histograms showing the percentage of streamlines to major fascicles for each of the functional regions in children (B) and adults (C); *Error bars:* standard error of the mean; *Dots*: individual participant data. (D,E) Correlation matrices depicting the average pairwise within-subject correlation between connectivity profiles for children (D) and adults (E). Lines depict cytoarchitectonic area; *cyan:* FG2, *magenta:* FG4.

In each subject, we quantified the fascicle connectivity profile for each functional ROI --that is, the percentage of streamlines of each functional ROI that connect to each of these five fascicles (IFOF, ILF, AF, pAF, and VOF). Consistent with the qualitative observations, ∼50% of the fascicle connections of mFus-faces and mOTS-words are to the pAF and 20-25% of the connections are to the ILF and AF, respectively (**Fig 3B**-top), whereas ∼50% of the fascicle connections of pFus-faces and pOTS-words are to VOF and ∼25% of the connections are to the pAF (**Fig 3B**-bottom). Similar fascicle connectivity profiles were observed in adults: ∼50% of the fascicle connections of mFus-faces and mOTS-words are to the pAF (**Fig 3C**-top) and ∼40-50% connections of pFus-faces and pOTS-words are to the VOF **(Fig 3C**-bottom).

Fascicle connectivity profiles shown in **Fig 3B,C** suggest that the fascicle connectivity profiles are similar among functional ROIs within the same cytoarchitectonic area consistent with the prediction that white matter is linked to cytoarchitecture (**Fig 2B**-Hypothesis 2). To quantify the similarity between connectivity profiles, we calculated the correlation between fascicle connectivity profiles of face- and word-selective regions within and across categories and cytoarchitectonic areas. If fascicle connectivity profiles are linked to category-selectivity, then functional ROIs selective for the same category would have connectivity profiles that are more correlated than functional ROIs located within the same cytoarchitectonic area. However, if fascicle connectivity profiles are linked to cytoarchitecture, then functional ROIs located within the same cytoarchitectonic area would have connectivity profiles that are more correlated than those selective for the same category.

The average within-subject correlation matrix for fascicle connectivity profiles is shown in **Fig 3D,E**. Results demonstrate that the correlations between connectivity profiles of functional ROIs within the same cytoarchitectonic area are higher than the correlations between functional ROIs with the same category-selectivity in both children (same cytoarchitectonic area: mOTS-words/mFus-faces correlation mean±sd: 0.85±0.19; pOTS-words/pFus-faces: 0.81±0.24; same category: mFus-faces/pFus-faces: 0.15±0.42; mOTS-words/pOTS-words: 0.27±0.46) and adults (same cytoarchitectonic area: mOTS-words/mFus-faces correlation mean±sd: 0.67±0.33; pOTS-words/pFus-faces: 0.77±0.30; same category: mFus-faces/pFus-faces: 0.26±0.49; mOTS-words/pOTS-words: 0.27±0.48). Similar results were observed (i) in the right hemisphere (**Supplemental Fig 13**), (ii) when controlling for motion and ROI size (**Supplemental Fig 14C**,**E)**, and (iii) for a restricted age range in children (**Supplemental Fig 14G**).

The significance of factors was evaluated using analyses-of-variance with Satterthwaite approximations for degrees of freedom (Kuznetsova et al., 2017) (referred to as linear mixed model analysis of variance, LMM ANOVA). A 2-way repeated-measures LMM ANOVA on the Fisher transformed correlation values between fascicle connectivity profiles with factors of shared feature (cytoarchitecture/category-selectivity) and age group (children/adults) revealed a significant main effect of shared feature (*F*(1,102.35) = 72.06, *p* = 1.72×10^−13^), reflecting the overall higher correlations between connectivity profiles located within the same cytoarchitectonic area compared to those with the same category-selectivity. There was no significant main effect of age group (*F*(1,42.34) = 1.34, *p* = 0.25), suggesting that correlations between connectivity profiles of the various functional ROIs were not different across children and adults. Finally, there was a significant age group by shared feature interaction (*F*(1,102.35) = 4.23, *p* = 0.04), reflecting higher correlations between connectivity profiles of functional ROIs located within the same cytoarchitectonic area compared to functional ROIs selective for the same category more so in children compared to adults.

### Decoupling cytoarchitecture and distance

One concern about these findings might be that functional ROIs located within the same cytoarchitectonic area may be physically closer than functional ROIs in different cytoarchitectonic areas. To test this possibility, we calculated the distances between functional ROIs within and across cytoarchitectonic areas in each subject and tested if there were systematic differences in distances between functional ROIs using a LMM repeated measures ANOVA with factors of cytoarchitecture area (same/different) and age group (children/adults). Functional ROIs within the same cytoarchitectonic areas were on average ∼10mm apart, but functional ROIs in different cytoarchitectonic areas were more than 15mm apart (**Fig 4A**-left). The LMM ANOVA revealed that indeed, functional ROIs in the same cytoarchitectonic area were closer than ROIs across cytoarchitectonic areas (main effect of cytoarchitecture: *F*(1,144) = 91.44, < 2×10^−16^). In general, the distance between ROIs did not differ between age groups (no significant effect of age group (*F*(1,144) = 0.71, *p* = 0.40). However, there was a significant group by cytoarchitecture interaction (*F*(1,144) = 6.43, *p* = 0.01), suggesting that ROIs within the same cytoarchitectonic areas are closer than ROIs in different cytoarchitectonic areas, but this difference is smaller in adults compared to children (**Fig 4A**-Functional ROI).

**Figure 4.**
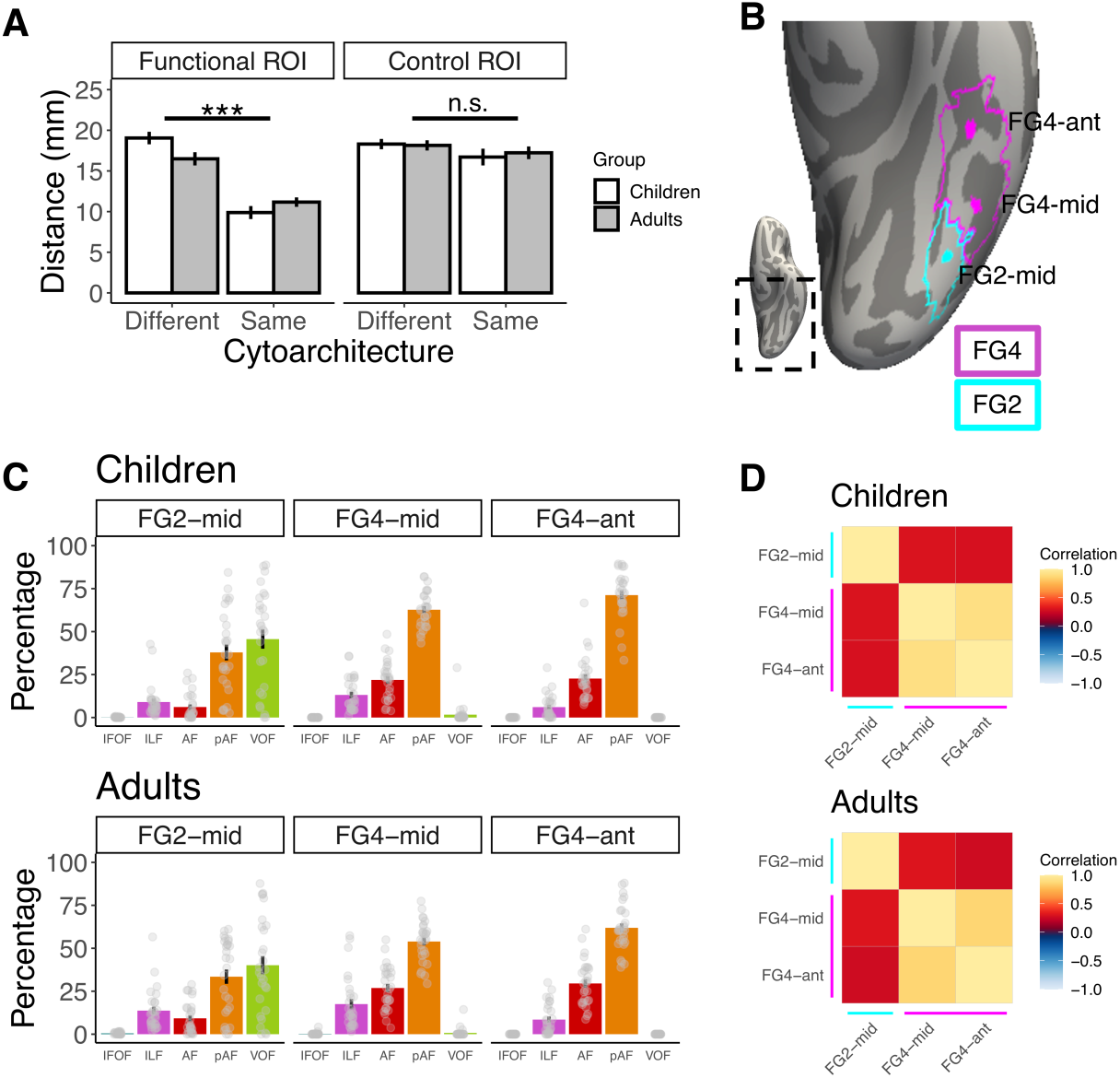
White matter is linked to cytoarchitecture when controlling for distance. **(A)** Average distance between functional ROIs located within the same and different cytoarchitectonic area. *Gray bars:* children, *white bars*: adults. *Error bars:* standard error of the mean. *Left:* average distance between functional ROIs; *Right:* average distance between control ROIs. LMM ANOVA on distances of the control ROI revealed no main effect of cytoarchitecture (*F*(1,106) = 2.56, *p* = 0.11), no main effect of age group (*F*(1,106) = 0.05, *p* = 0.82) and no interaction (*F*(1,106) = 0.19, *p* = 0.66)). *Significance*: *** p < .001. (B) Inflated fsaverage cortical surface depicting control ROIs. Colors indicate cytoarchitectonic area: *cyan:* FG2, *magenta:* FG4. *Acronyms: FG4-mid:* control region in mid FG4. *FG4-ant:* control region in anterior FG4, *FG2-mid:* control region in mid FG2. Lines indicate cytoarchitectonic area: *cyan:* FG2; *magenta:* FG4. (C) Fascicle connectivity profiles for each of the equally spaced disk ROIs in children (top) and adults (bottom); *Error bars:* standard error of the mean; *Dots*: individual participant data. *Acronyms: IFOF:* inferior fronto-occipital fasciculus; *ILF:* inferior longitudinal fasciculus; *AF:* arcuate fasciculus; *pAF:* posterior arcuate fasciculus; *VOF:* vertical occipital fasciculus. (D) Correlation matrices depicting the average within-subject correlation between fascicle connectivity profiles of equally spaced ROIs in children (top) and adults (bottom).

To assess if spatial proximity is driving the results, we conducted two control analyses (i) we added distance as factor to the LMM ANOVA, and (ii) we examined if equidistant ROIs within and across cytoarchitectonic areas had the same or different fascicle connectivity.

Results of the first analysis adding distance as a covariate to the LMM ANOVA revealed that even as distance among ROIs contributes to the similarity among their fascicle connectivity profiles (main effect of distance: *F*(1,139.71) = 19.81, *p* = 1.73×10^−5^), connectivity profiles of ROIs located within the same cytoarchitectonic area were significantly more correlated than those selective for the same category (main effect of shared feature: *F*(1,118.79) = 16.50, *p* = 8.77×10^−5^). There were no significant effects of group (*F*(1,43.60) = 2.40, *p* = 0.13) or a shared feature by group interaction (*F*(1,105.66) = 1.39, *p* = 0.24).

For the second analysis, we compared the functionally defined white matter for equidistant control ROIs (**Fig 4B**), two of which were in FG4 (FG4-mid/FG4-ant) and one was in FG2 (FG2-mid); the middle ROI (FG4-mid) was located equally far from FG4-ant and FG2-mid along a posterior-to-anterior axis (**Fig 4A**-control ROI). We reasoned that if distance drives the results, then the connectivity profile of FG4-mid would be equally correlated to that of FG4-ant and FG2-mid because they are equally far apart, consistent with the predictions of the posterior-anterior connectivity gradient hypothesis. However, if the correlation between FG4-mid and FG4-ant is significantly higher than between FG4-mid and FG2-mid, then it would provide strong evidence that white matter connectivity profiles are coupled with cytoarchitecture, not distance.

Results in **Fig 4C** reveal that in both children and adults, connectivity profiles of ROIs that are equally spaced apart and located in the same cytoarchitectonic area (FG4-mid and FG4-ant) are more similar than those across cytoarchitectonic areas (FG4-mid and FG2-mid). FG4-mid and FG4-ant are connected primarily to the pAF whereas FG2-mid connects to both the VOF and pAF. Indeed, pairwise correlations between connectivity profiles are significantly higher among equally spaced ROIs within the same cytoarchitectonic area than across cytoarchitectonic areas (**Fig 4D**, significant main effect of cytoarchitecture (*F*(1,53) = 75.84, *p* = 8.41×10^−12^), no significant differences across age groups (*F*(1,53) = 0.77, *p* = 0.38), and no group by cytoarchitecture interaction (*F*(1,53) = 0.78, *p* = 0.38, LMM ANOVA on the pairwise Fisher transformed correlations)). These results indicate that connectivity profiles are linked to cytoarchitecture independent of the distance between the ROIs.

### Do fascicle connectivity profiles of VTC ROIs predict cytoarchitecture or category-selectivity?

We further hypothesized that if there is a consistent relationship between fascicle connectivity profiles and cytoarchitecture or between fascicle connectivity profiles and category, it should be possible to predict the cytoarchitectonic area or category-selectivity, respectively, from an unlabeled fascicle connectivity profile. We tested these hypotheses using a leave-one-out maximum correlation classification approach. That is, in each iteration, we calculated the average connectivity profile of each cytoarchitectonic area (FG2/FG4) and category (faces/words) using data from n-1 subjects. Then, we performed two separate two-way classifications of the left-out subject’s connectivity profiles: (i) the cytoarchitectonic area (FG2 or FG4), and (ii) the category-selectivity (face or word).

For children (**Fig 5**, light colors), the classifier successfully predicted the cytoarchitectonic area, but not category selectivity above chance (chance level: 50%; cytoarchitecture: mean classification accuracy±sd = 82%±38%; *t*(78) = 7.47, *p* = 1.01×10^−10^; category-selectivity: mean classification accuracy±sd = 49%±50%; *t*(78) = -0.11, *p* = 0.91). For adults (**Fig 5**, dark colors), the classifier successfully predicted both cytoarchitectonic area and category-selectivity above chance (cytoarchitecture: mean classification accuracy±sd = 87%±34%; *t*(96) = 10.53, *p* = 2.2×10^−16^; category: mean classification accuracy±sd = 66%±48%; *t*(96) = 3.31, *p* = 0.001). Correct classification was significantly higher for cytoarchitecture vs. category (odds ratio = 3.32, 95% CI [1.62, 6.84], binominal logistic regression with factors of classification type and age group) and significantly lower for children vs. adults (odds ratio = .50, 95% CI [0.27,0.92]), but there was no significant interaction (odds ratio: 1.43, 95% CI [0.51, 3.97]). Together, these analyses provide strong evidence supporting the hypothesis that fascicle connectivity profiles better predict cytoarchitecture rather than category-selectivity across age groups.

**Figure 5.**
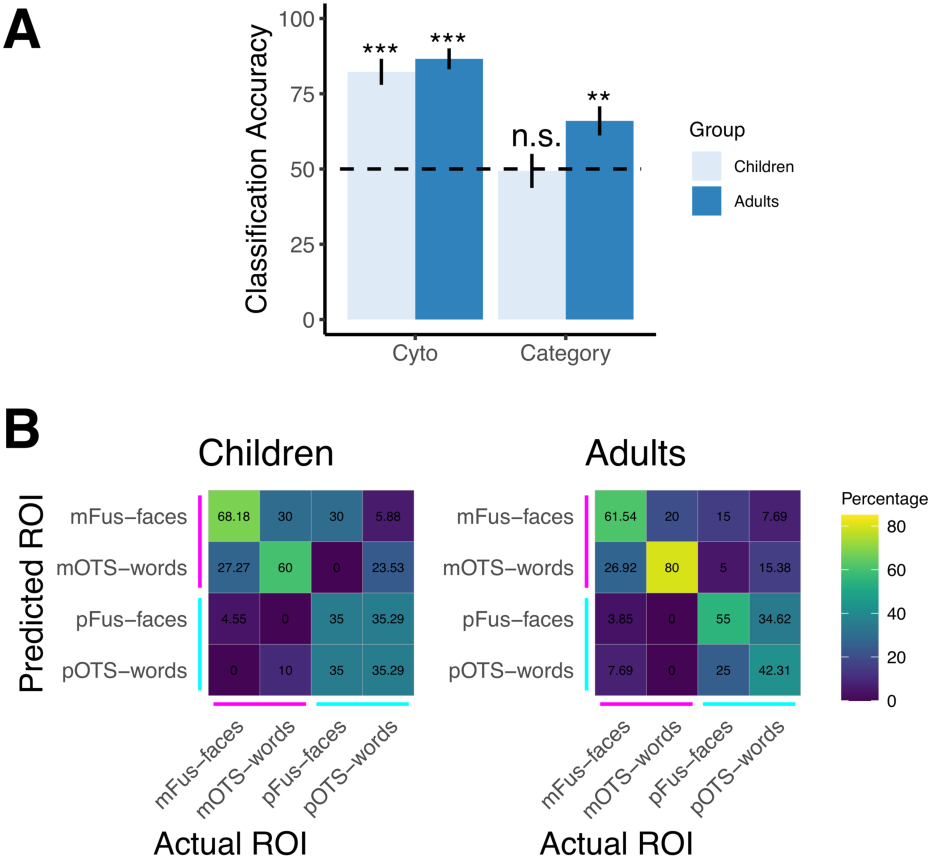
Fascicle connectivity profiles predict cytoarchitecture more than category. **(A)** Average classification across participants of either the cytoarchitectonic area or category-selectivity from the fascicle connectivity profiles. *Light blue:* Children, dark blue: adults; *Error bars:* standard error of the mean. Dashed line: chance level. Significance vs. chance: * p < .05, ** p < .01, *** p < .001. n.s.: not significant. (B) Confusion matrices for the average classification and misclassification of functionals ROI across participants. Each cell depicts the percentage of times each actual functional ROI (x-axis) was predicted (y-axis), see colorbar. *On-diagonals:* % successful classification; *off-diagonals:* % miss-classification. Each column sums to 100%. *Acronyms:* mFus-faces: mid fusiform face-selective region. pFus-faces: posterior fusiform face-selective region. mOTS-words: mid occipitotemporal sulcus word-selective region. pOTS-words: posterior occipitotemporal sulcus word-selective region. *Cytoarchitectonic areas:* cyan: FG2, magenta: FG4.

Next, we examined the patterns of misclassification on a four-way ROI classification to see whether ROIs were more often misclassified as ROIs located within the same cytoarchitectonic area or selective for the same category. Confusion matrices in **Fig 5B** confirm that for both children and adults, ROIs were more often misclassified as ROIs located within the same cytoarchitectonic area. This result confirms that connectivity profiles were more similar for ROIs located within the same cytoarchitectonic area. For three out of the four ROIs, classification (on-diagonal values in confusion matrices) improved from childhood to adulthood, suggesting that connectivity profiles of specific ROIs become more distinguishable over development.

### Endpoint connectivity profiles of VTC functional ROIs are organized by cytoarchitecture

While the fascicle connectivity profile reveals which fascicles connect to each functional ROI, it is also important to understand the endpoint connectivity profiles of the white matter connections of each functional ROI. The endpoint connectivity profile provides complementary data to the fascicle connectivity profile for three reasons: (i) Not all fibers in the whole brain connectome are part of major fascicles, e.g., many fibers connecting VTC to early visual cortex are not classified as part of the major fascicles in common atlases (Catani et al., 2002; Wakana et al., 2004). (ii) Endpoint profiles provide additional information as to which cortical regions VTC functional ROIs connect to, and (iii) Functional ROIs may share fascicles but have different endpoints, e.g., mOTS-words and mFus-faces may both connect to the arcuate fasciculus, but only mOTS-words may have endpoints in Broca’s area.

Therefore, we next examined the endpoint connectivity profiles of the VTC functional ROIs. To do so, we intersected each of the face- and word-selective functional ROIs with the whole brain connectome of each subject. To determine the cortical regions to which these white matter tracts connect, we projected the tract endpoints to the cortical surface, and then used the Glasser Atlas (Glasser et al., 2016) to quantify the distribution of the endpoints across the brain.

**Fig 6** shows the average spatial distribution across cortex of white matter endpoints for each of the VTC face and word-selective functional ROIs in children (**Fig 6A**) and adults (**Fig 6B**). Qualitatively, the spatial distribution of endpoints is similar between anterior functional ROIs (mFus-faces and mOTS-words) as well as similar between posterior functional ROIs (pFus-faces and pOTS-words). The anterior functional ROIs (mFus-faces and mOTS-words) show a distinct spatial distribution of endpoints with the highest endpoint density in ventral occipital and anterior temporal cortex, followed by endpoints in parietal cortex and inferior frontal cortex. In contrast, the posterior functional ROIs (pFus-faces and pOTS-words) illustrate the highest endpoint density in ventral temporal and occipital cortex, followed by the posterior parietal cortex and anterior temporal cortex. In other words, the endpoint density for posterior face- and word-selective regions appears to be shifted more posteriorly compared to that of anterior face- and word-selective regions.

**Figure 6.**
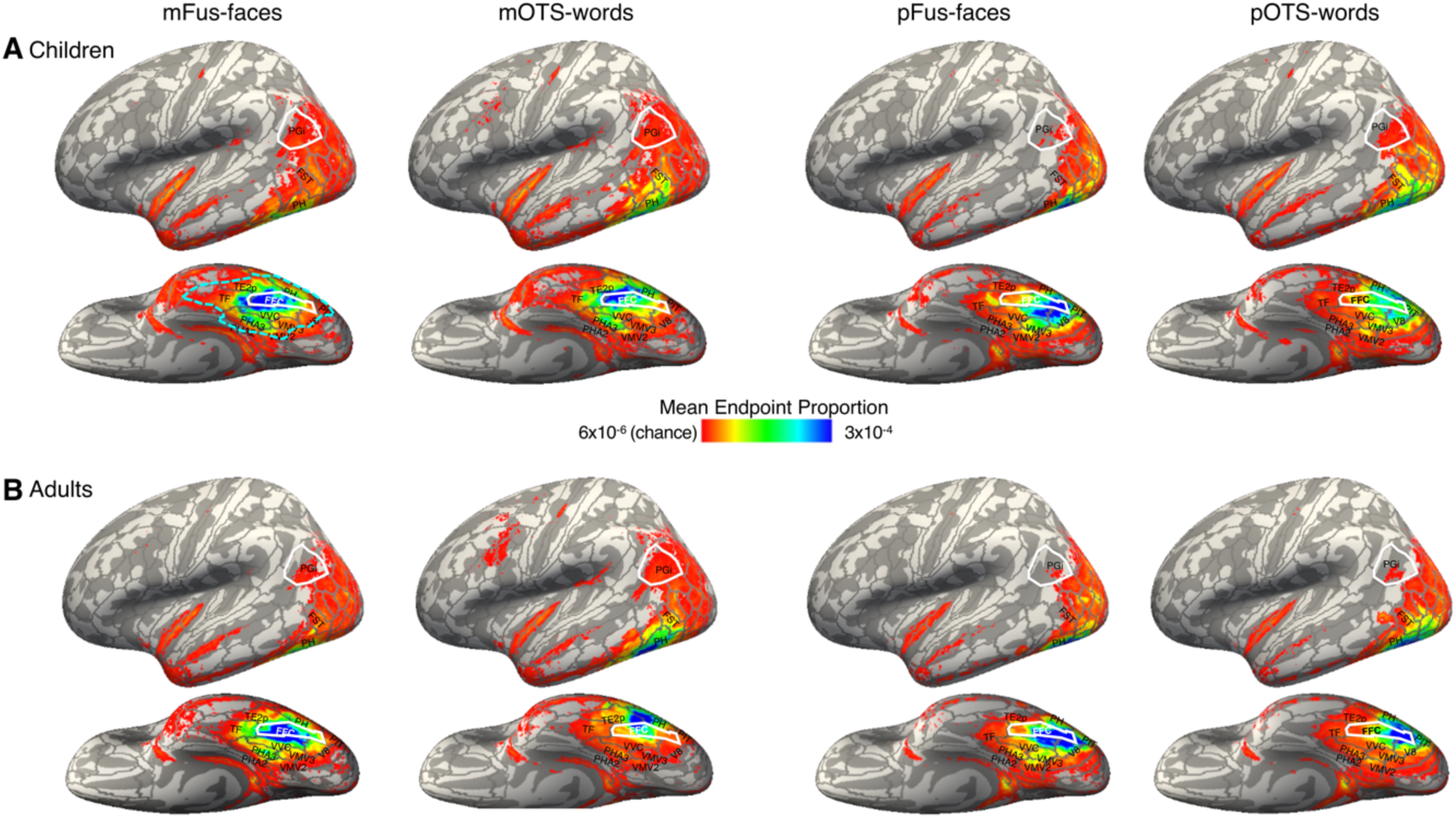
Endpoint connectivity profile of face- and word-selective regions across the entire brain. Average endpoint connectivity profile of mFus-face, pFus-faces, mOTS-words, pOTS-words in children (A) and adults (B). Data are shown on the fsaverage inflated cortical surface. *Color map:* endpoint proportion at each vertex (each map sums to one). *Threshold value of maps:* chance level if all endpoints were evenly distributed across the brain. *Gray outlines:* Glasser ROIs. *Cyan outline*: ventral temporal cortex. *White outlines:* PGi and FFC as a visual reference to aid comparison across panels.

We next quantified the endpoint distributions of each of the functional ROIs across the brain using the Glasser Atlas parcellation in children (**Fig 7A**) and adults (**Fig 7B**). Right hemisphere data is in **Supplemental Fig 15**. Consistent with the qualitative analysis, the endpoint distributions for ROIs located in the same cytoarchitectonic area are similar. That is, pFus-faces and pOTS-words have similar endpoint distributions, with the highest endpoint densities within the occipital lobe, and mFus-faces and mOTS-words have similar endpoint distributions, with higher endpoint proportions in the temporal and occipital lobes than the parietal and frontal lobes. Interestingly, mFus-faces and mOTS-words appear to have more distinct endpoint distributions in adults (**Fig 7B**) compared to children (**Fig 7A**), especially in the occipital lobe.

**Figure 7.**
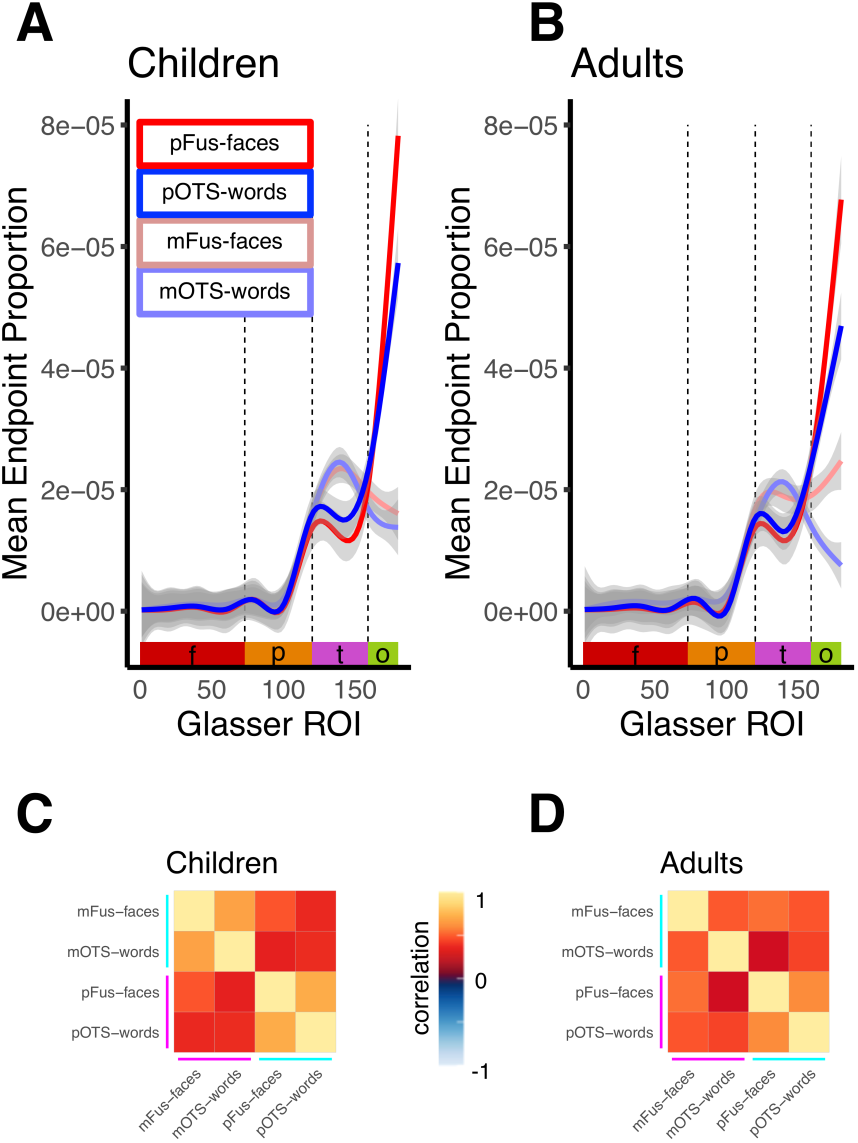
Quantification of endpoint connectivity profile of face- and word-selective regions in children and adults. **(A,B)** Mean endpoint proportion across 180 Glasser ROIs for each of the VTC word and face-selective regions in children (A) and adults (B). Line color indicates the functional ROI; *red:* pFus-faces; *blue:* pOTS-words; *light red:* mFus-faces; *light blue:* mOTS-words. *Shaded area:* standard error of the mean. *X-axis:* Glasser Atlas ROI number arranged by lobe: *red:* frontal (f); *orange:* parietal (p); *magenta:* temporal (t); *green:* occipital (o). *Vertical dashed lines:* lobe boundaries. (C,D) Correlation matrices depicting the average within-subject pairwise correlation between endpoint connectivity profiles of ventral face and word-selective ROIs in children (C) and adults (D).

To quantitatively test our hypotheses (**Fig 2B**), we calculated the correlation between the endpoint density profiles of all pairs of functional ROIs, and then tested if correlations between endpoint profiles were higher among functional ROIs within the same cytoarchitectonic area or among functional ROIs selective for the same category. The correlation matrix in **Fig 7C** demonstrates that for children, functional ROIs within the same cytoarchitectonic area had higher correlations among endpoint profiles compared to functional ROIs selective for the same category (mean correlation±sd, same cytoarchitectonic area: mOTS-words/mFus-faces: 0.68±0.24; pOTS-words/pFus-faces: 0.71±0.20; same category: mFus-faces/pFus-faces: 0.48±0.19; mOTS-words/pOTS-words: 0.38±0.16). For adults (**Fig 7D**), however, this pattern was less clear (same cytoarchitectonic area: mFus-faces/mOTS-words 0.49±0.24, pFus-faces/pOTS-words 0.60±0.24; same category: mFus-faces/pFus-faces: 0.54±0.22, mOTS-words/pOTS-words: 0.44±0.21).

A 2-way repeated-measures LMM ANOVA on Fisher transformed correlations between connectivity profiles with factors of shared feature (cytoarchitecture/category-selectivity) and age group (children/adults), with distance as an independent factor, revealed that even as distance contributes to the correlations among endpoint connectivity profiles (*F*(1,138.91) = 36.78, *p* = 1.18×10^−8^), endpoint connectivity profiles are overall more correlated in children than adults (main effect of age group: *F*(1, 36.41) = 5.54, *p* = 0.02). Additionally, endpoint connectivity profiles are significantly more correlated among functional ROIs within the same cytoarchitectonic area than functional ROIs with the same category-selectivity in children compared to adults (feature by age group interaction *F*(1,98.43) = 5.24, *p* = 0.02; post-hoc t-test, cyto vs. category, children: *t*(39.93) = 4.70, *p* = 3.08×10^−5^; adults: *t*(78.72) = 1.23, *p* = 0.22). In other words, we find evidence for significant developmental differences in endpoint connectivity profiles. In children, but not adults, the endpoint connectivity profiles are more correlated among functional ROIs within the same cytoarchitectonic area compared to functional ROIs selective for the same category.

As we found a feature by age interaction and prior literature suggests that the spatial extent of face and word-selective functional ROIs increases from childhood to adulthood (Turkeltaub et al. 2003; Scherf et al. 2007; Golarai et al. 2010; Nordt et al. 2021), we asked if these age group differences were due to developmental changes to the functional ROIs. Thus, we repeated endpoint profile analyses in two ways (i) controlling for functional ROI size (**Supplemental Fig 16C**), and (ii) excluding connections to the Glasser ROIs in VTC (**Supplemental Figs 16E, 17**). We reasoned that if group differences are eliminated under these controls, it would suggest that developmental changes in VTC are driving age group differences in connectivity profiles. Alternatively, finding a similar pattern of results in the control analyses would provide strong evidence for development differences in endpoint connectivity profiles that supersede local changes in the VTC. Both control analyses replicate the findings in **Fig 7**. That is, we find significantly higher correlations among endpoint connectivity profiles of functional ROIs within the same cytoarchitectonic areas than among functional ROIs of the same category preference and significantly more so in children than adults (**Supplemental Fig 16A**,**C**,**E**,**G**,**I)**. Additional controls of matching motion across children and adults (**Supplemental Fig 16G)** and analysis of a subset of children in a more restricted age range (**Supplemental Fig 16I**) also replicate these findings.

### Endpoint connectivity profiles better predict cytoarchitecture than category in children, but not adults

To further examine the predictive nature of the endpoint connectivity profiles, we used a leave-one-out maximum correlation classification approach to test whether we can predict cytoarchitectonic area (FG2/FG4) or category-selectivity (faces/words) from an unlabeled endpoint connectivity profile. The classifier successfully predicted cytoarchitecture above chance for both children (**Fig 8A**-light colors, mean classification accuracy±sd: 90%±30%; *t*(78) = 11.67, *p* = 2.2×10^−16^) and adults (**Fig 8A**-dark colors, mean accuracy±sd: 84%±37%; *t*(96) = 8.84, *p* = 4.49×10^−14^). Additionally, the classifier predicted category-selectivity significantly above chance for both children (mean accuracy±sd: 70%±46%; *t*(78) = 3.77, *p* = 0.0003) and adults (mean accuracy±sd: 80%±40%; *t*(96) = 7.51, *p* = 3.07×10^−11^). There were no significant differences in the odds of correct classification of cytoarchitecture compared to category (odds ratio = 1.23, 95% CI [0.59, 2.57], binominal logistic regression), or for children compared to adults (odds ratio = 0.56, 95% CI [0.28,1.12], binominal logistic regression). There was, however, a significant interactive effect of classification type by age group (odds ratio: 3.14, 95% CI [1.00, 9.83], binominal logistic regression), reflecting higher classification accuracy of cytoarchitecture than category in children but not adults. Results were largely similar when controlling for functional ROI size, excluding Glasser ROIs in VTC, matching for motion, and using a restricted age range for the child subjects (**Supplemental Fig 16B**,**D**,**F**,**H**,**J)**. These data suggest that in childhood, endpoint connectivity profiles predict cytoarchitecture better than category-selectivity, but by adulthood, this is no longer the case.

**Figure 8.**
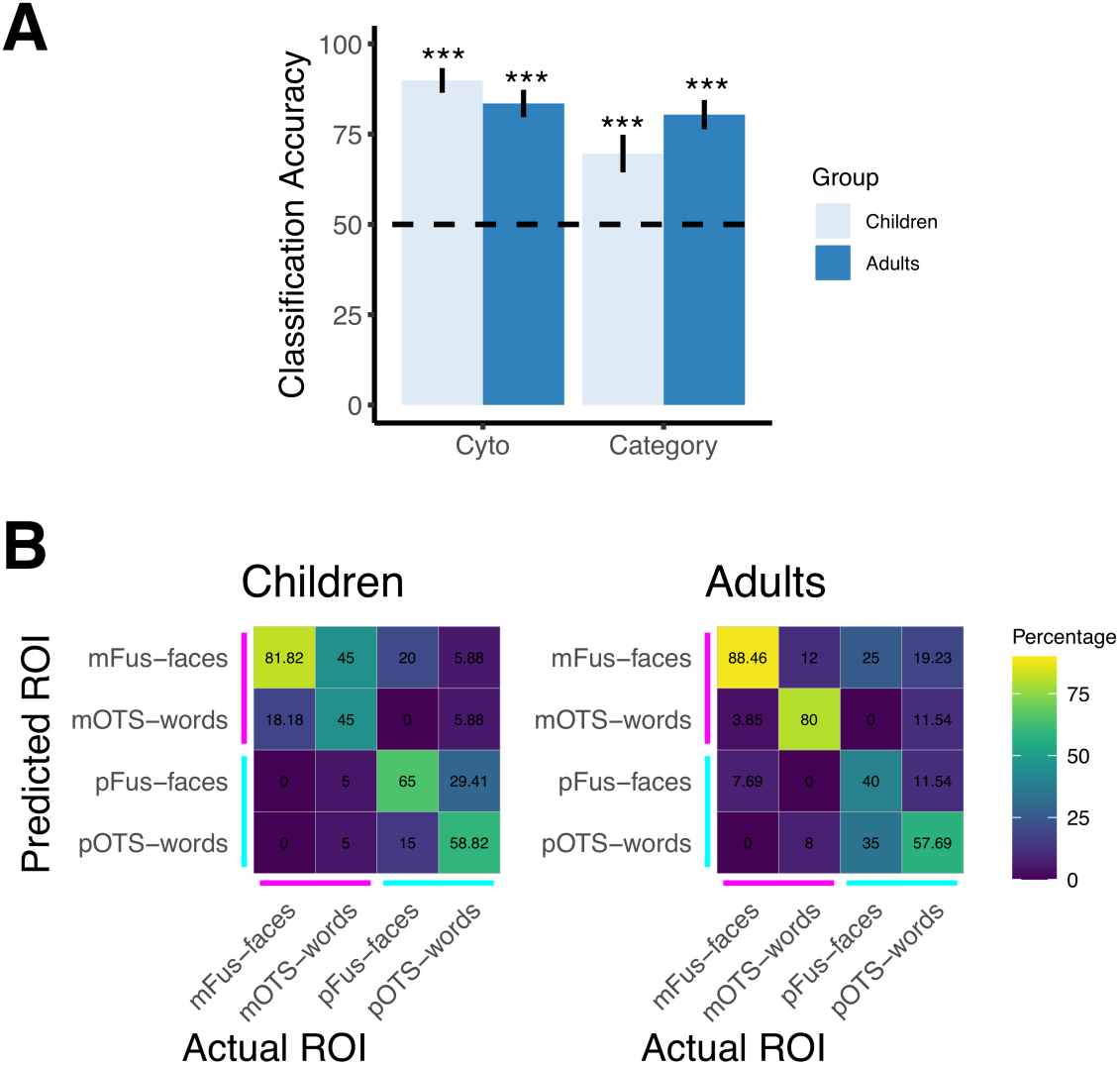
Endpoint connectivity profiles better predict cytoarchitecture than category in children, but not adults. **(A)** Bar graphs showing classification accuracy of either the cytoarchitectonic area or category-selectivity from the endpoint connectivity profiles. *Light blue:* Children, dark blue: adults; *Error bars:* standard error of the mean. *Dashed line:* chance level. Significantly different than chance: * p < .05, ** p < .01, *** p < .001. *Acronyms*: mFus-faces: mid fusiform face-selective region. pFus-faces: posterior fusiform face-selective region. mOTS-words: mid occipitotemporal sulcus word-selective region. pOTS-words: posterior occipitotemporal sulcus word-selective region. (B) Average classification and misclassification of functional ROI across participants. Each cell depicts the percentage of times each actual functional ROI (x-axis) was predicted (y-axis), see colorbar. On-diagonals: % successful classification; off-diagonals: % misclassification. *Acronyms:* mFus-faces: mid fusiform face-selective region. pFus-faces: posterior fusiform face-selective region. mOTS-words: mid occipitotemporal sulcus word-selective region. pOTS-words: posterior occipitotemporal sulcus word-selective region. *Cytoarchitectonic areas*: cyan: FG2, magenta: FG4.

To understand which ROIs drive the increased category classification for adults compared to children, we implemented a four-way classifier that predicts the functional ROI associated with an unlabeled connectivity profile in left out subjects, and then examined the pattern of errors. The classification confusion matrix (**Fig 8B**) indicates that anterior functional ROIs (mFus-faces and mOTS-words) were better classified in adults than children, but the opposite was true for posterior functional ROIs (pFus-faces and pOTS-words). Interestingly, the largest developmental increase in accuracy was observed for mOTS-words, as classification of mOTS-words from endpoint connectivity profiles almost doubled from children to adults, resulting from more distinct connectivity profiles for mOTS-words and mFus-faces in adults compared to children.

Overall, the endpoint connectivity profile analysis of ventral functional ROIs suggests that initially in childhood there is a tight link between endpoint connectivity profiles and cytoarchitecture, as endpoint connectivity predicts cytoarchitecture better than category-selectivity and confusability is largely among functional ROIs within the same cytoarchitectonic area. However, development from childhood to adulthood in the endpoint connectivity of anterior functional ROIs, and particularly mOTS-words, results in more distinct endpoint connectivity profiles for the functional ROIs within FG4, suggesting that in adulthood both cytoarchitecture and category-selectivity contribute to connectivity profiles.

### Distinguishing cytoarchitecture from a posterior-to-anterior gradient

Here, we find converging evidence that connectivity profiles are organized by cytoarchitecture rather than category-selectivity in childhood. However, the multiple word and face selective regions are organized along a posterior-to-anterior anatomical axis that is thought to be linked to a hierarchical processing axis. This observation raises the question of whether the connectivity profiles of the face and word ROIs reflect the posterior-anterior hierarchical axis of these functional ROIs or is specific to their cytoarchitecture.

To address this possibility, we leveraged the other category-selective regions in VTC. OTS-bodies is a region that is selective for bodies and located in cytoarchitectonic area FG4 (together with mFus-faces and mOTS-words), whereas CoS-places is a region selective for scenes and is located in cytoarchitectonic area FG3 in both children and adults (Weiner et al., 2017; **Supplementary Fig 18**). Additionally, along this posterior-to-anterior axis, CoS-places and OTS-bodies are similarly anterior as mFus-faces and mOTS-words. Considering the connectivity profiles for these additional functional ROIs would therefore allow us to dissociate the two hypotheses. We reasoned that if the connectivity profiles are organized by cytoarchitecture, then we would expect that OTS-bodies would have a similar connectivity profile to mFus-faces and mOTS-words and that CoS-places would have a distinct connectivity profile from all the other ROIs (**Fig 9A**). However, if connectivity profiles reflect differences along a posterior-to-anterior axis, then we would expect that CoS-places and OTS-bodies would both have similar connectivity profiles to mFus-faces and mOTS-words as they are all more anterior ROIs (**Fig 9B**). Note that both hypotheses would still predict that the connectivity profiles of these four anterior functional ROIs would be different than the posterior ones (pFus-faces, pOTS-words).

**Figure 9.**
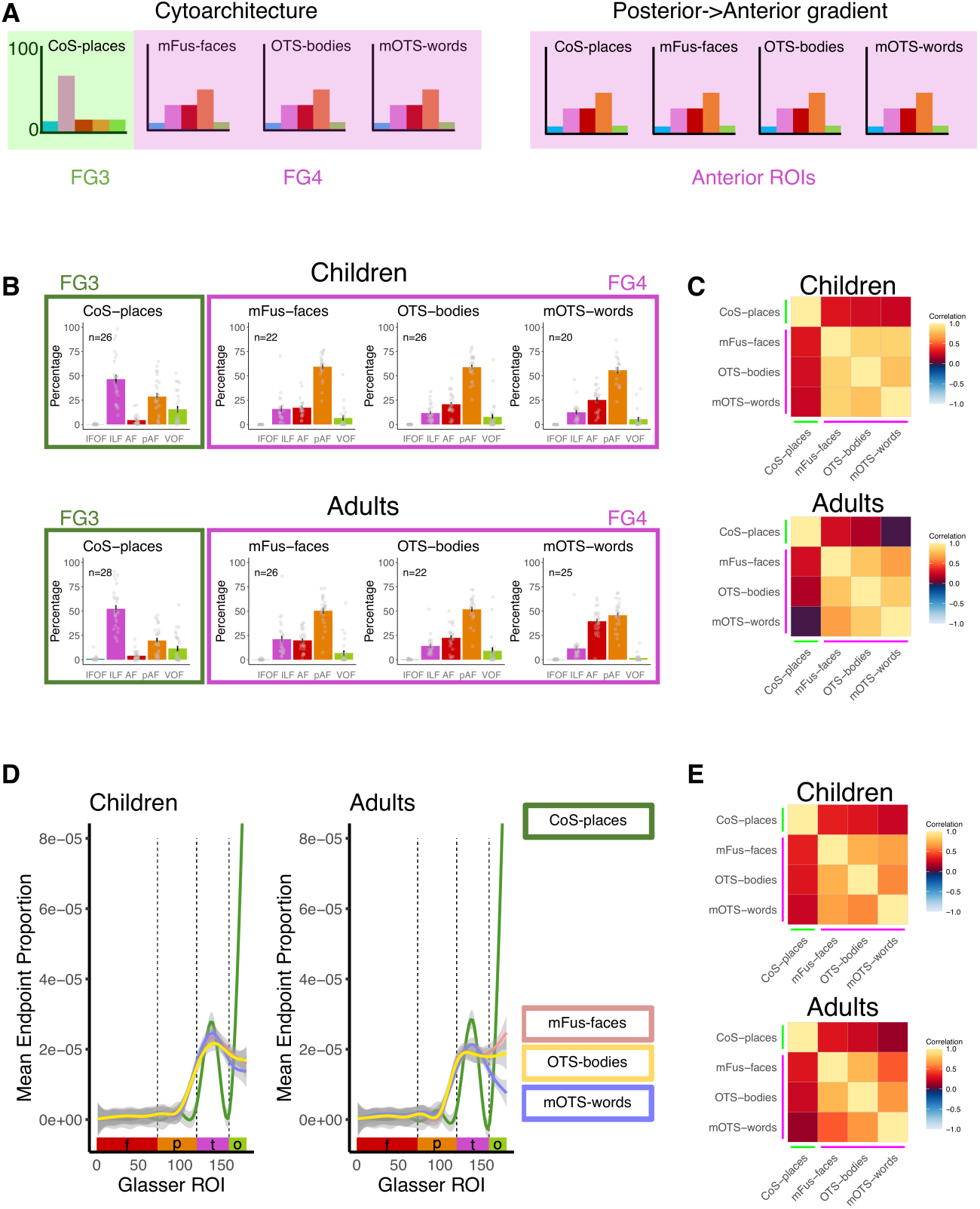
Distinguishing cytoarchitecture from a posterior-to-anterior gradient. **(A)** Schematic illustration of the predicted fascicle connectivity profile of anterior category-selective ROIs according to the cytoarchitectonic (left) and posterior-to-anterior hypotheses). Each panel shows the predicted distributions of connections across white matter fascicles for a given functional ROI. *Bar Colors*: fascicles. CoS-places: collateral sulcus place-selective region; mFus-faces: mid fusiform face-selective region; mOTS-words: mid occipito-temporal sulcus word-selective region. OTS-bodies: occipito-temporal body selective region. (B) *Left:* fascicle connectivity profile of these functional ROIs in children and adults. *Error bars:* standard error of the mean; *Dots:* individual participant data. *Right:* Correlation matrices depicting the average pairwise within-subject correlation between connectivity profiles of these four functional ROIs for children (top) and adults (bottom). Lines depict cytoarchitectonic area; *green:* FG3, *magenta*: FG4. (**C**) *Left:* endpoint connectivity profile of these functional ROIs across 180 Glasser ROIs in children (left) and adults (right). Line color indicates the functional ROI: *green:* CoS-places; *yellow:* OTS-bodies; *light red:* mFus-faces; *light blue:* mOTS-words; *Shaded area:* standard error of the mean. X-axis: Glasser Atlas ROI number arranged by lobe: *red:* frontal (f); *orange:* parietal (p); *magenta:* temporal (t); *green:* occipital (o). *Vertical lines:* lobe boundaries. *Right*: Correlation matrices depicting the average pairwise within-subject correlation between endpoint connectivity profiles for children (top) and adults (bottom). Lines depict cytoarchitectonic area; *green:* FG3, *magenta:* FG4.

**Fig 9B-left** shows the fascicle connectivity profiles for these four category-selective regions in children and adults. In both age groups, the majority of the fascicle connections of OTS-bodies are to the pAF followed by the AF and ILF. This fascicle connectivity profile is similar to that of mFus-faces and mOTS-words. In contrast, CoS-places has a distinct connectivity profile with the majority of its fascicle connections going to the ILF followed by the pAF and VOF. Quantitative analyses of the pairwise correlations between endpoint fascicle connectivity profiles (**Fig 9B-right**) show that the connectivity profile of the CoS-places is dissimilar from that of the other three category-selective regions (main effect of cytoarchitecture, *F*(1,467.17) = 463.72, *p* < 2×10^−16^, LMM ANOVA on Fisher transformed correlation values). Additionally, fascicle connectivity profiles were more similar in children than adults (main effect of age group: F(1,46.79) = 5.05, p = 0.03; no significant interaction F(1,467.17) = 1.86, p = 0.17).

Similar results are found for the endpoint connectivity profiles, where mFus-faces, mOTS-words, and OTS-bodies have similar connectivity profiles and CoS-place has a distinct connectivity profile with many more connections to the occipital lobe as compared to the other three ROIs (**Fig 9C-left**). Quantitative analyses of the pairwise correlations among endpoint connectivity profiles (**Fig 9C-right)** show a significant effect of cytoarchitecture (*F*(1,474.89) = 369.81, *p* < 2×10^−16^), as well a significant main effect of age group (*F*(1,47.35) = 4.28, *p* = 0.04 but no group by cytoarchitecture interaction (*F*(1,474.89) = 0.27, *p* = 0.60). Overall, these results suggest that connectivity profiles are linked to cytoarchitecture rather than a posterior-to-anterior gradient along VTC.

## Discussion

Here, we tested whether white matter connections of category-selective functional ROIs in the ventral stream are linked to category-selectivity or cytoarchitecture. We find that in both children and adults, fascicle connectivity profiles better predict cytoarchitecture than category-selectivity. In adulthood, however, endpoint connectivity profiles for functional ROIs located within the same cytoarchitectonic region became more distinct, particularly in FG4. These results suggest that in childhood, cytoarchitecture and white matter largely co-vary together. However, over development connectivity profiles may become more specialized to support category-specific processing, resulting in connectivity profiles linked both to cytoarchitecture and category-selectivity.

### The relationship between functional regions and cytoarchitecture is consistent across children and adults

First, we measured the relationship between cytoarchitecture and functional ROIs in children and adults. Importantly, we replicate Weiner et al., (2017) and find that both in children and adults there is a consistent relationship between cytoarchitecture and category-selective regions, where mFus-faces and mOTS-words are located in cytoarchitectonic area FG4, and pFus-faces and pOTS-words are located in cytoarchitectonic area FG2. These results suggest that functional ROIs in VTC have a consistent organization across development. It is important to note that the cytoarchitectonic areas were estimated from adult brains (Amunts et al. 2000; Rottschy et al. 2007; Caspers et al. 2013; Lorenz et al. 2017), and due to the paucity of pediatric postmortem tissue, no histological study to date has compared the cytoarchitecture of VTC in children and adults. Future histological research can test whether cytoarchitecture is consistent over childhood development, though several lines of evidence suggest that childhood development of VTC tissue likely involves increases in myelination and dendritic arborization rather than changes in cell density (Gomez et al. 2017; Natu et al. 2019).

### Rethinking the relationship between white matter and category-selectivity

After establishing the relationship between cytoarchitectonic areas and category selective regions, we were able to test whether white matter is linked to cytoarchitectonic area or category selectivity. Prior work has suggested that category-selective regions have specific white matter tracts to support category-selective processing (Mahon and Caramazza 2011; Bouhali et al. 2014; Li et al. 2020). Some of the most compelling work demonstrating the relationship between category-selectivity and white matter connectivity are fingerprinting studies that have shown that it is possible to predict category-selective responses from the endpoint connectivity profile across the brain (Saygin et al. 2011; Osher et al. 2016; Grotheer, Yeatman, et al. 2021), and further that it is possible to predict from endpoint connectivity at age five where word-selective activations will emerge in VTC at age eight (Saygin et al. 2016). In this study, Saygin et al., (2016) found that the correlation between the predicted word-selectivity and actual word selectivity from the white matter fingerprint is 0.47 ± 0.036 (mean across subjects, Fisher z). It is interesting that we find that the correlation between connectivity profiles of functional ROIs selective for the same category is comparable (children: 0.49 ± 0.04, adults: 0.59±0.05, mean Fisher z). Strikingly, however, the correlation between connectivity profiles for functional ROIs located in the same cytoarchitectonic area is much higher (children:1.03±0.11, adults:0.69±0.06, mean Fisher z). We also predict category-selectivity from white matter connectivity above chance for both children and adults. However, by testing predictivity of white matter for multiple attributes including ones that have not been tested before, we find that connectivity profiles better predict cytoarchitecture than category-selectivity.

It is important to note that some category-selective regions that were tested in prior studies are also located in different cytoarchitectonic areas. For example, face-selective and place-selective regions have both unique cytoarchitecture (Weiner et al. 2017) and unique white matter connectivity (Saygin et al. 2011; Gomez et al. 2015). Therefore, comparing the white matter connectivity profiles of face- and place-selective regions may lead to the inaccurate conclusion that category-selectivity, cytoarchitecture, and white-matter are all linked. Instead, our study shows that this is not the general case, and it is necessary to examine cases where category-selective and cytoarchitecture are orthogonal to reveal the broader principle that white matter is linked to cytoarchitecture above category-selectivity.

### Developmental Implications

The theory that white matter connections are linked to category-selectivity also has developmental implications. For example, it has been proposed that white matter early in childhood provides scaffolding for category-selectivity and that innate white matter connections may explain similarities in the functional organization of visual cortex between blind and sighted subjects (Mahon and Caramazza 2011; Bi et al. 2016; van den Hurk et al. 2017; Murty et al. 2020).

Here, we find that early in childhood white matter is better tied to cytoarchitecture than category, and for the fascicle connectivity profiles the link between cytoarchitecture and white matter is consistent over development. In contrast, the endpoint connectivity profiles better predict cytoarchitecture than category in childhood, but not in adulthood. These findings suggest a rethinking of the view that white matter tracts provide an innate scaffold for category-selective function. Instead, it seems that white matter and cytoarchitecture are closely linked early in development, and that category-specific connections can emerge over development. Recent findings reveal that white matter fascicles can be reliably identified in individual newborns (Grotheer et al. 2022) paving the way for future research to examine whether the relationship between cytoarchitecture and white matter connections is present at birth.

An interesting direction for future research is understanding the mechanisms that enable tuning of white matter connections during childhood development as well as developmental changes to the properties of the white matter itself (Yeatman, Dougherty, Ben-Shachar, et al. 2012; Yeatman, Wandell, et al. 2014; Moulton et al. 2019; Vinci-Booher et al. 2022). Importantly, we measure the white matter connections that connect to category-selective regions. Therefore, differences in white matter connections could reflect changes in the white matter or changes in functional selectivity over development. If the white matter is changing, a mechanism underlying development may be the pruning or growth of connections to support the more specific connections needed for brain networks to better support specific behaviors. On the other hand, as functional regions in VTC develop and grow (Turkeltaub et al. 2003; Scherf et al. 2007; Golarai et al. 2010; Nordt et al. 2021), they may shift to the edges of cytoarchitectonic areas, thus, forming unique connections that span multiple networks. For example, examination of the probabilistic maps of mOTS-words in children and adults suggest that while it is located lateral to the mid fusiform sulcus in both children and adults, its location is more variable in children compared to adults (**Fig 1B**). Therefore, more distinct connectivity profiles for mOTS-words in adults could be associated the mOTS-words shifting laterally from childhood to adulthood, potentially becoming more aligned with the endpoints of the arcuate fasciculus. Since our data suggest that white matter connections are domain general early in development, one implication from this study is that interventions targeting certain skills (e.g., reading) may be more successful before white matter connections gain category-specificity.

### What is the utility of shared connectivity?

While prior work has argued for the utility of specific white matter connections to support domain specific processing, there are also reasons why it might be useful to have white matter connections that are domain general. Similar white matter connections of functional regions within a cytoarchitectonic area suggest that regions within the same cytoarchitectonic may perform similar computations. An example of such a shared computation across the domains of reading and face recognition is the processing of foveal information which requires high visual acuity (Levy et al. 2001; Hasson et al. 2002).

An additional benefit of shared white matter connections for functional regions selective for different categories may be flexibility. A large body of research has revealed that learning can lead to the development of new category-selectivity in VTC. For example, illiterate children and adults do not have word-selective regions in VTC; however, both groups develop word-selectivity after learning to read (Brem et al. 2010; Dehaene et al. 2010; Saygin et al. 2016). A flexible and domain-general system may enable new selectivity to emerge as it becomes relevant, as well as older selectivity (e.g., selectivity to limbs) to be recycled (Nordt et al. 2021). Since it is not possible to predict which categories will be behaviorally relevant (e.g., children can learn artificial categories such as Pokémon (Gomez et al. 2019)), a domain-general architecture provides flexibility to develop many category representations.

In addition to shared computational demands for regions selective for different categories, regions in different cytoarchitectonic areas may be performing different computations on the same stimulus. For example, classic studies suggested that unique cytoarchitecture and white matter connections enable parallel processing of different attributes of a stimulus (such as its motion and color) (Zeki and Shipp 1988; Van Essen et al. 1992). In the context of VTC, prior research has demonstrated that word-selective regions located within different cytoarchitectonic regions perform different computations on the same stimuli (Lerma-Usabiaga et al. 2018; White et al. 2019; Caffarra et al. 2021). These studies have shown that (i) pOTS-words can process two words in parallel, whereas mOTS-words can only process one word at a time (White et al. 2019) and (ii) pOTS-words is involved in extracting visual features, whereas mOTS-words is involved in processing lexical features (Lerma-Usabiaga et al. 2018). Therefore, we propose that multiple regions that are selective for the same category but located in different cytoarchitectonic regions and have different white matter connections may perform different computations towards a shared visual behavior.

### Beyond VTC

In addition to elucidating the pattern of white matter connections of the human visual system, this work has broader implications for the rest of the brain. Specifically, we hypothesize that across the brain, there are tight links between white matter connections and cytoarchitecture. Other evidence supporting this idea comes from past work showing that supplementary motor area (SMA) and pre-SMA which are in distinct cytoarchitectonic regions also have distinct connectivity profiles (Johansen-Berg et al. 2004; Klein et al. 2007), as well as studies that showed that Broca’s area spans multiple cytoarchitectonic regions (Amunts et al. 2010) with distinct connectivity profiles (Klein et al. 2007). Future research can examine this hypothesis in other sensory-motor (Shipp 2003; Heil et al. 2005; Morosan et al. 2005) and cognitive systems (Sporns et al. 2005; Siegel et al. 2012) across the brain. In addition, future research can examine how white matter connectivity profiles and cytoarchitecture relate to other features of cortex such as gene expression, neurotransmitter receptor distribution (Markello et al. 2022) and cortical folding (Kaneko et al. 2020).

### Summary

In school-aged children and adults the white matter connections of ventral functional ROIs are linked to cytoarchitecture. Over development, white matter connectivity profiles become more distinct for functional ROIs with different category selectivity located within the same cytoarchitectonic region. These findings suggest that cytoarchitecture and white matter are linked early in childhood and become more specific during development. Together our data propose a rethinking of the view that category-selective regions have innate and specific white matter connections to support category-specific processing.

## Acknowledgments

We thank Brianna Jeska and Michael Barnett for their contributions in scanning subjects.

