## Supplemental Fig for "White matter connections of high-level visual areas predict cytoarchitecture better than category-selectivity in childhood, but not adulthood"

Supplemental Figures

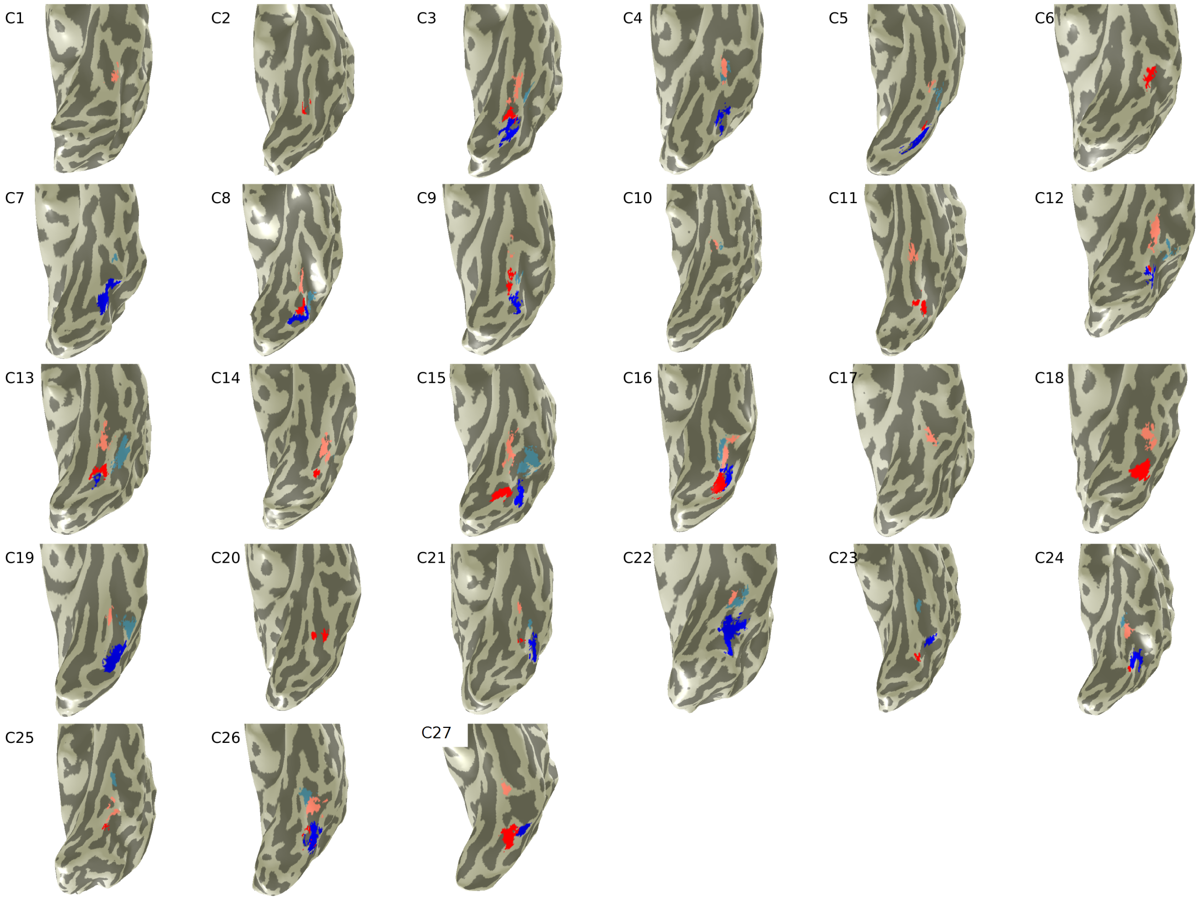


**Supplemental Figure 1. Functional regions of interest (ROIs) in the left hemisphere of individual child participants (ordered by age).** Upper left of each image: *C:* child participant, *number:* child's ranking youngest to oldest. *Light blue:* mOTS-words, *dark blue:* pOTS-words, *pink:* mFus-faces, *red:* pFus-faces.

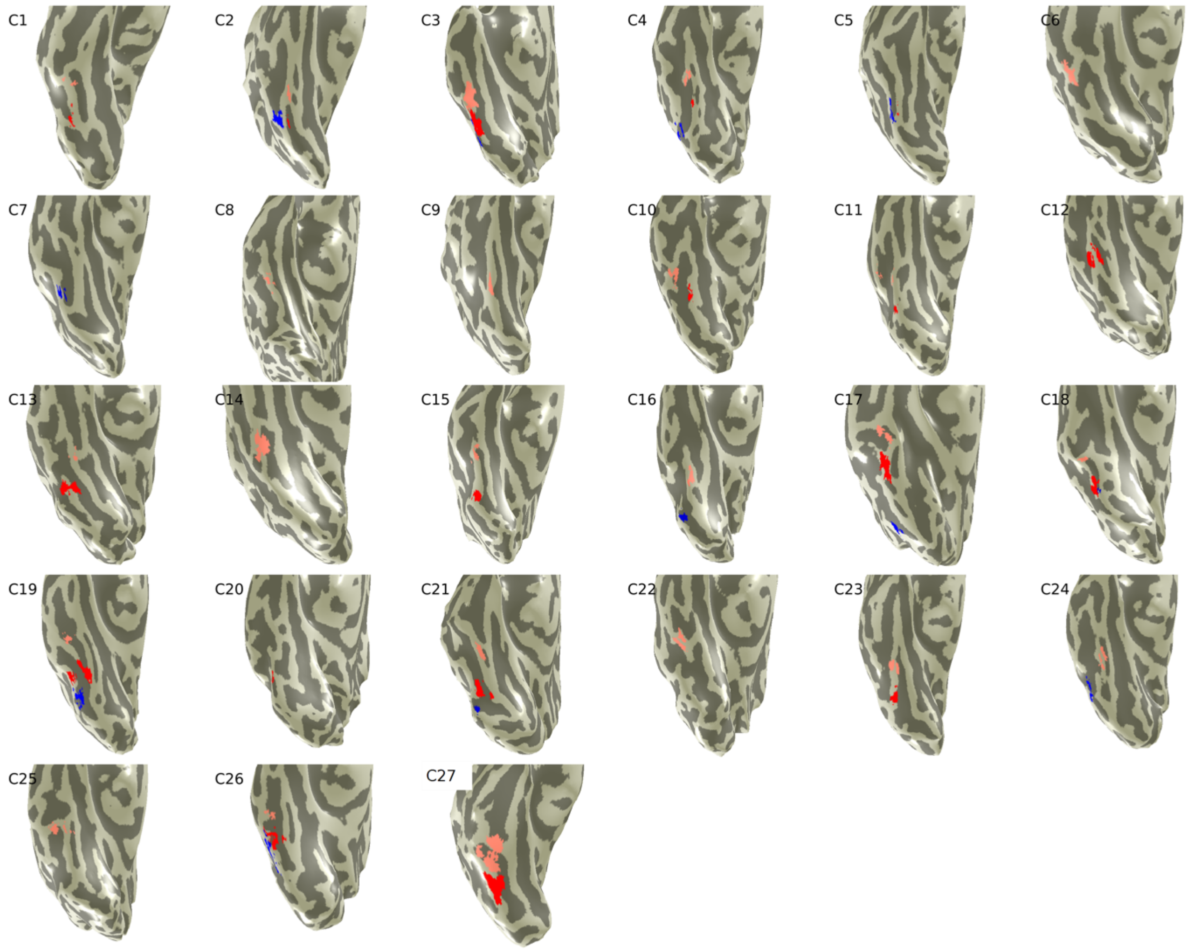


**Supplemental Figure 2. Functional ROIs in the right hemisphere of individual child participants (ordered by age).** Upper left of each image: *C:* child participant, *number:* child's ranking youngest to oldest. *Light blue:* mOTS-words, *dark blue:* pOTS-words, *pink:* mFus-faces, *red:* pFus-faces.

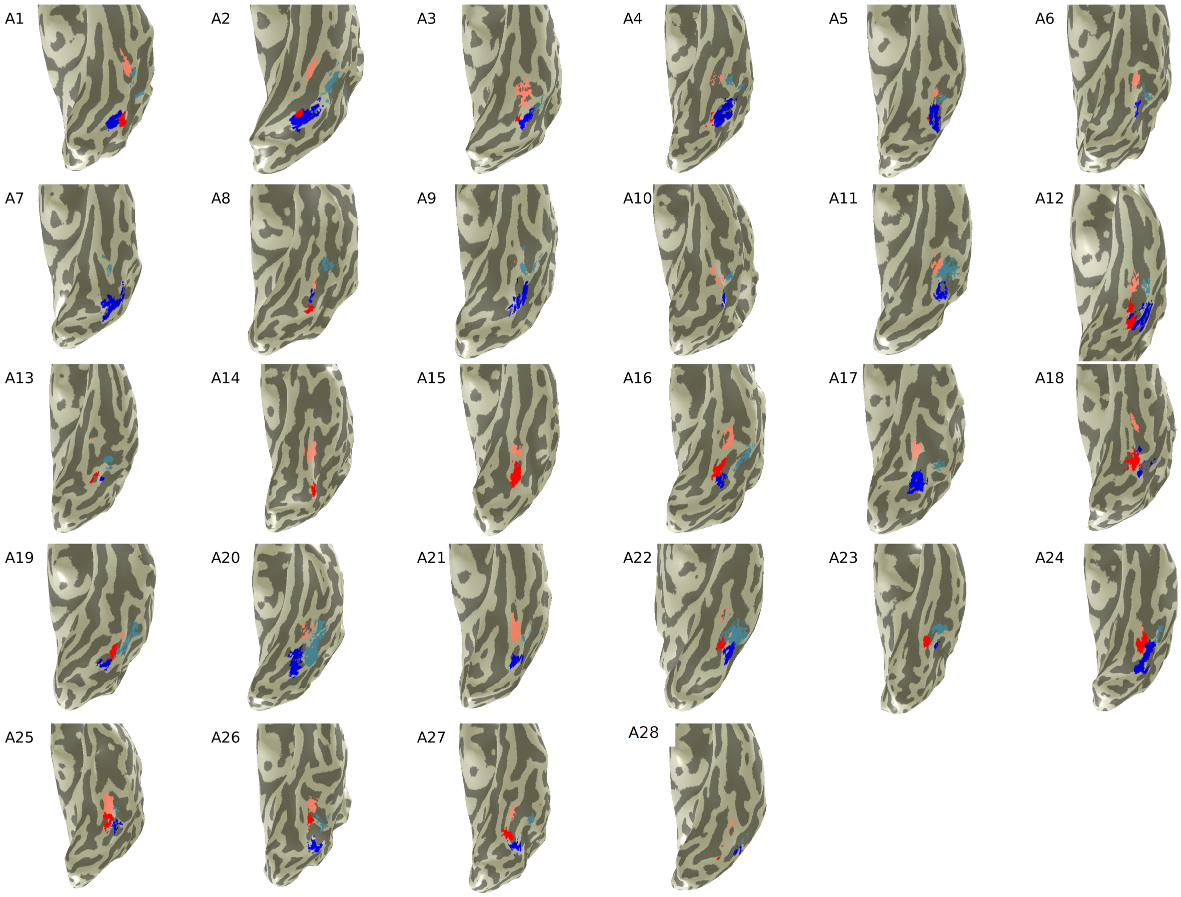


**Supplemental Figure 3. Functional ROIs in the left hemisphere of individual adult participants (ordered by age).** Upper left of each image: *A:* adult participant, *number:* adult's ranking youngest to oldest. *Light blue:* mOTS-words, *dark blue:* pOTS-words, *pink:* mFus-faces, *red:* pFus-faces.

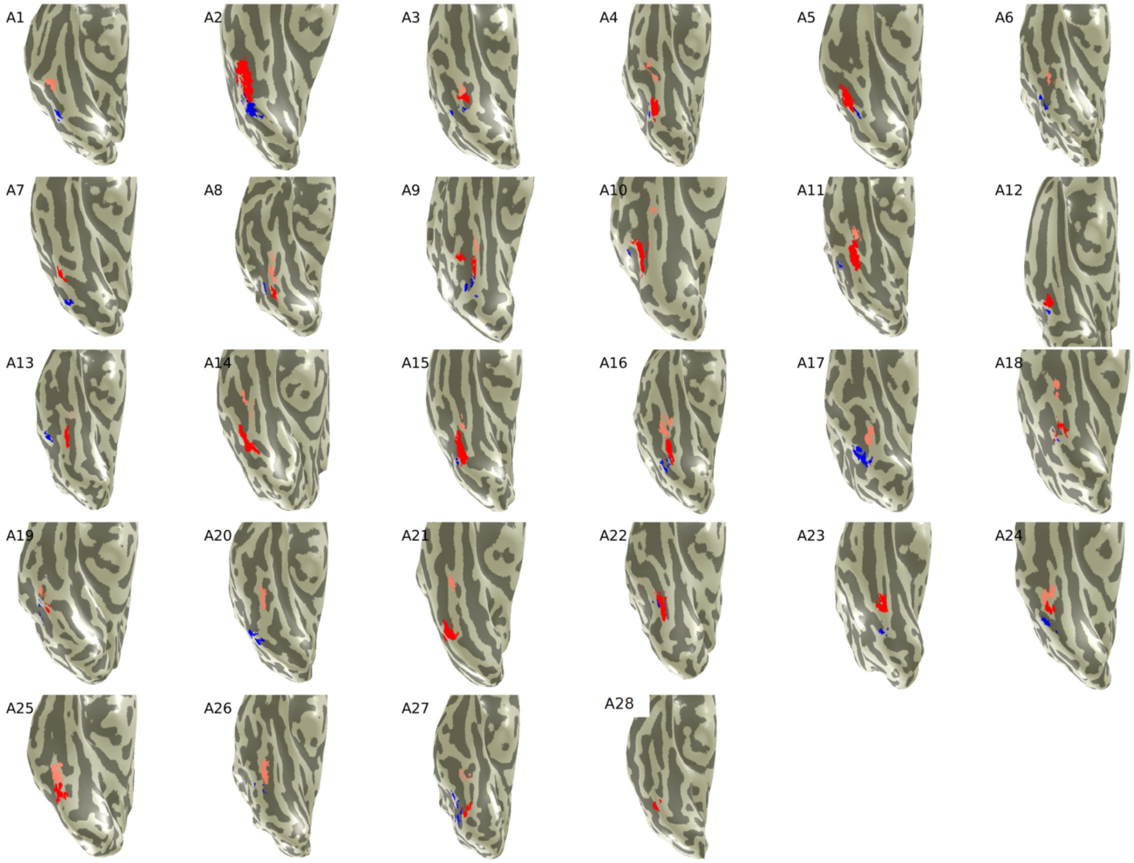


**Supplemental Figure 4. Functional ROIs in the right hemisphere of adult participants (ordered by age) in the right hemisphere.** Upper left of each image: *A:* adult participant, *number:* adult's ranking youngest to oldest. *Light blue:* mOTS-words, *dark blue:* pOTS-words, *pink:* mFus-faces, *red:* pFus-faces.

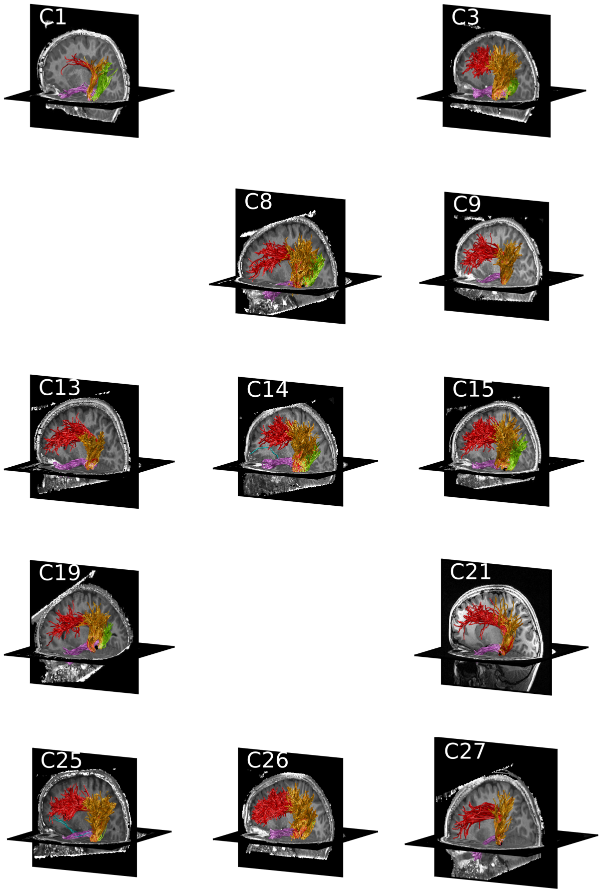


**Supplemental Figure 5. Functionally defined white matter tracts for mFus-faces in individual child participants (ordered by age).** Upper left of each image: *C:* child participant, *number:* child's ranking youngest to oldest. *Black:* functional ROI: mFus-faces. Fascicles: *blue:* inferior frontal occipital fasciculus, *magenta:* inferior longitudinal fasciculus, *red:* arcuate fasciculus, *orange:* posterior arcuate fasciculus, *green:* vertical occipital fasciculus.

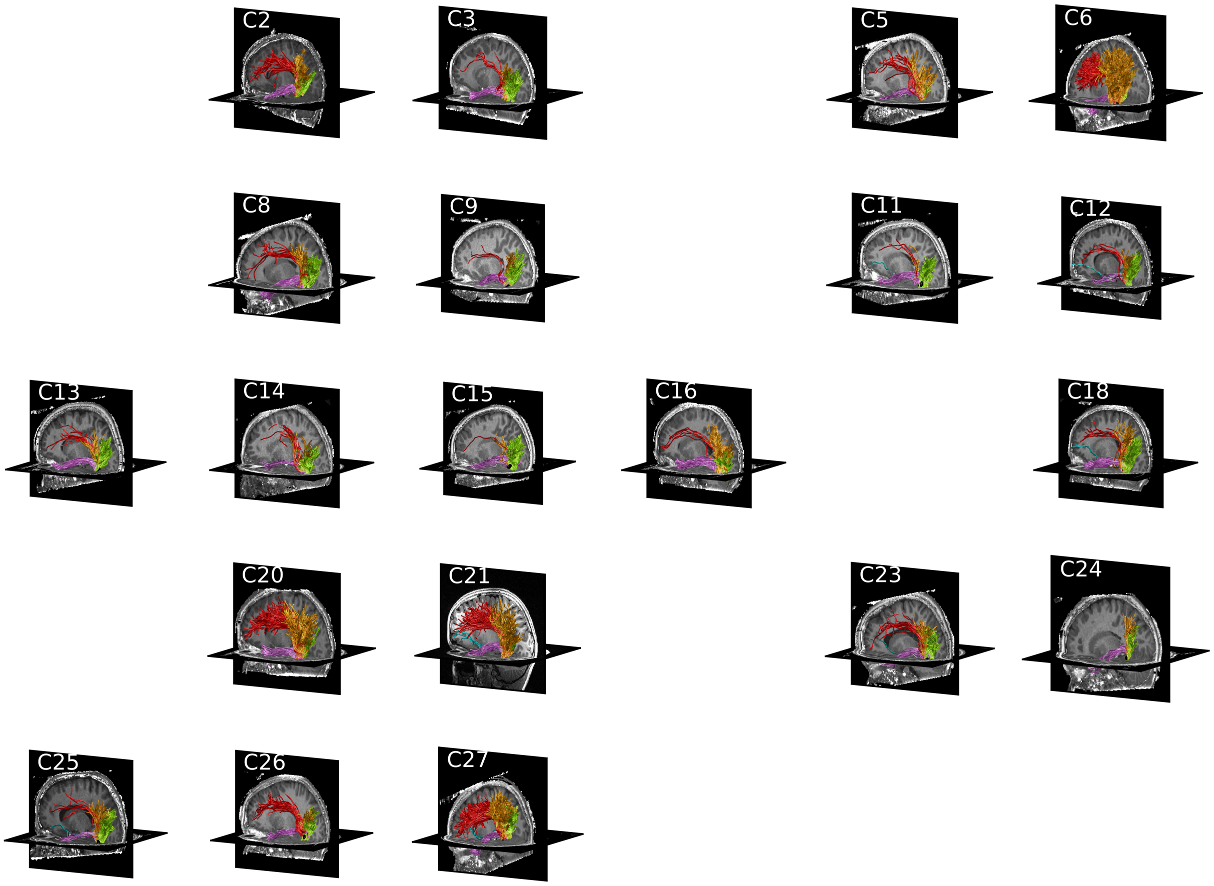


**Supplemental Figure 6. Functionally defined white matter tracts for pFus-faces in individual child participants (ordered by age).** Upper left of each image: *C:* child participant, *number:* child's ranking youngest to oldest. *Black:* functional ROI: pFus-faces. Fascicles: *blue:* inferior frontal occipital fasciculus, *magenta:* inferior longitudinal fasciculus, *red:* arcuate fasciculus, *orange:* posterior arcuate fasciculus, *green:* vertical occipital fasciculus.

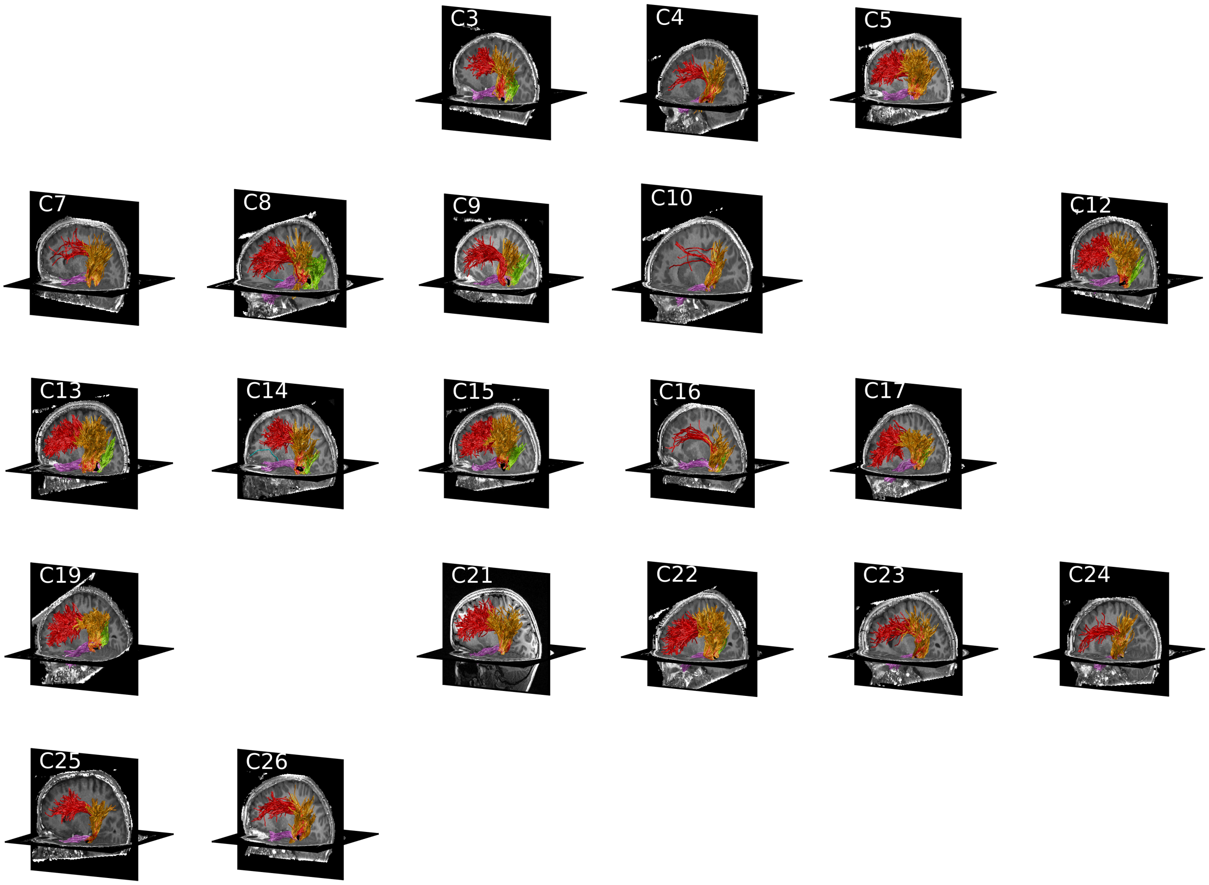


**Supplemental Figure 7. Functionally defined white matter tracts for mOTS-words in individual child participants (ordered by age).** Upper left of each image: *C:* child participant, *number:* child's ranking youngest to oldest. *Black:* functional ROI: mOTS-words. Fascicles: *blue:* inferior frontal occipital fasciculus, *magenta:* inferior longitudinal fasciculus, *red:* arcuate fasciculus, *orange:* posterior arcuate fasciculus, *green:* vertical occipital fasciculus.

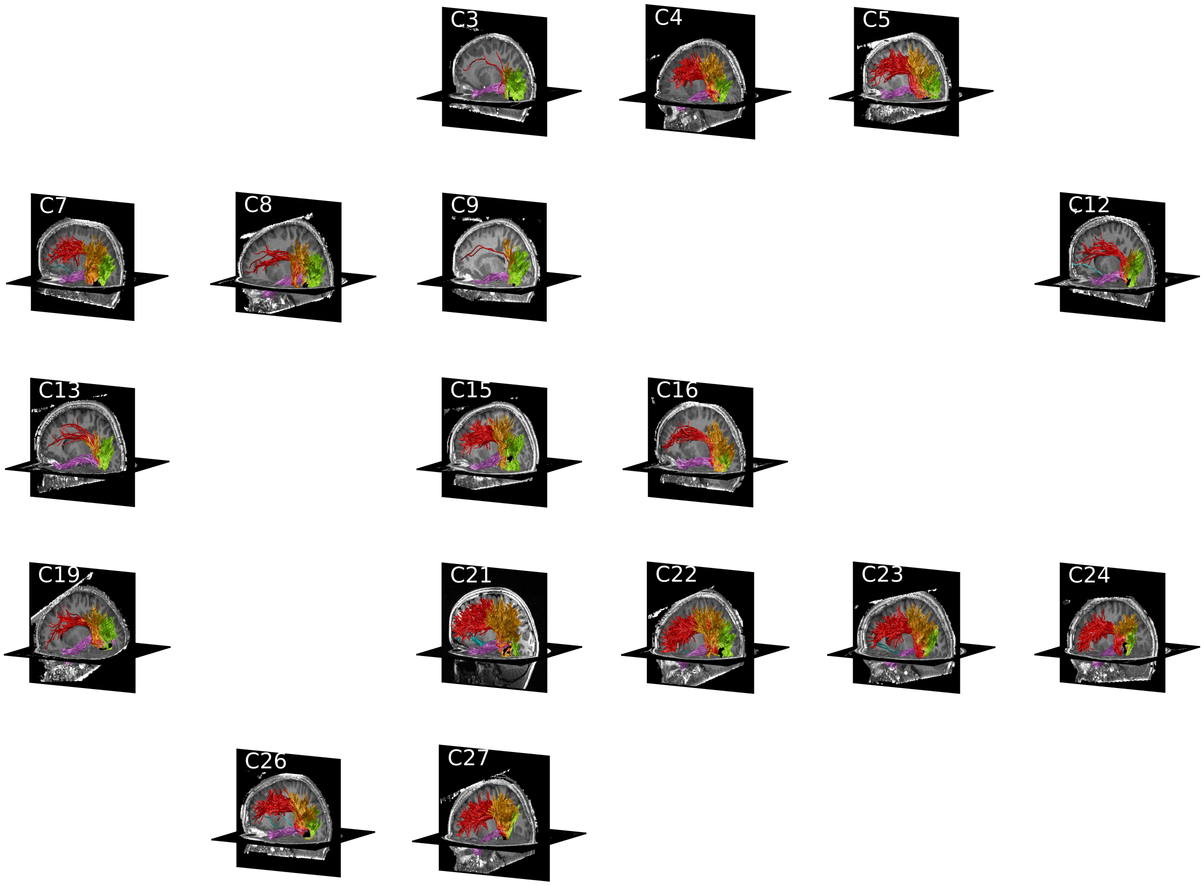


**Supplemental Figure 8. Functionally defined white matter tracts for pOTS-words in individual child participants (ordered by age).** Upper left of each image: *C:* child participant, *number:* child's ranking youngest to oldest. *Black:* functional ROI: pOTS-words. Fascicles: *blue:* inferior frontal occipital fasciculus, *magenta:* inferior longitudinal fasciculus, *red:* arcuate fasciculus, *orange:* posterior arcuate fasciculus, *green:* vertical occipital fasciculus.

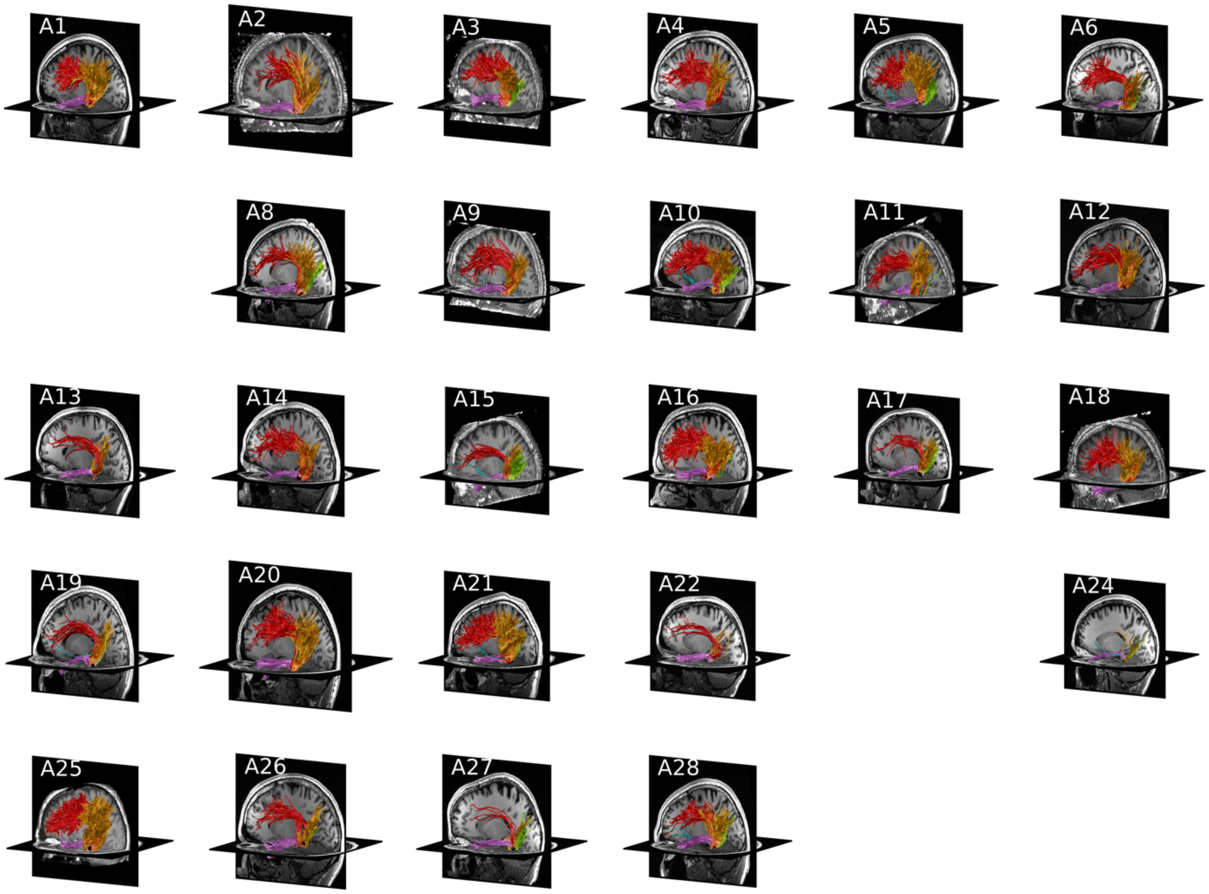


**Supplemental Figure 9. Functionally defined white matter tracts for mFus-faces in individual adult participants (ordered by age).** Upper left of each image: A*:* adult participant, *number:* adult's ranking youngest to oldest. *Black:* functional ROI: mFus-faces. Fascicles: *blue:* inferior frontal occipital fasciculus, *magenta:* inferior longitudinal fasciculus, *red:* arcuate fasciculus, *orange:* posterior arcuate fasciculus, *green:* vertical occipital fasciculus.

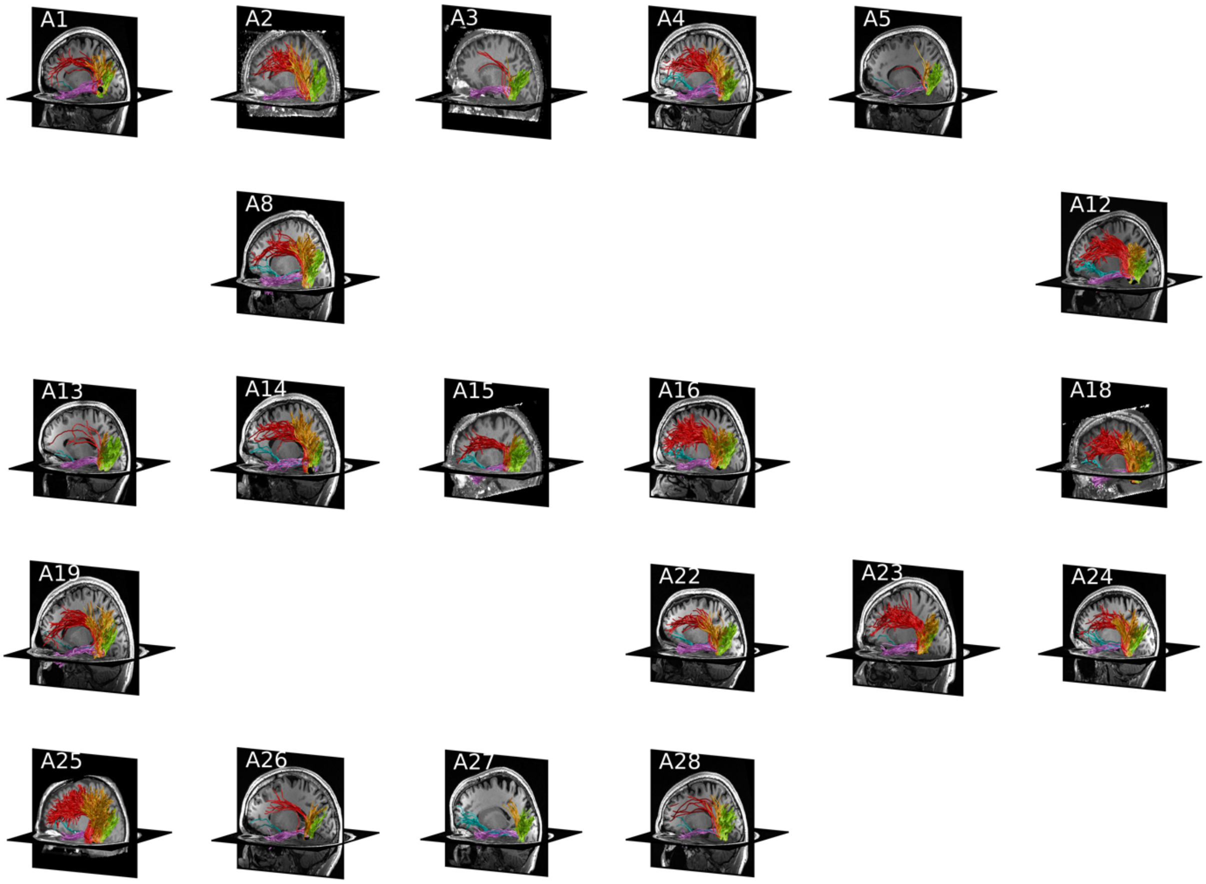


**Supplemental Figure 10. Functionally defined white matter tracts for pFus-faces in individual adult participants (ordered by age).** Upper left of each image: A*:* adult participant, *number:* adult's ranking youngest to oldest. *Black:* functional ROI: pFus-faces. Fascicles: *blue:* inferior frontal occipital fasciculus, *magenta:* inferior longitudinal fasciculus, *red:* arcuate fasciculus, *orange:* posterior arcuate fasciculus, *green:* vertical occipital fasciculus.

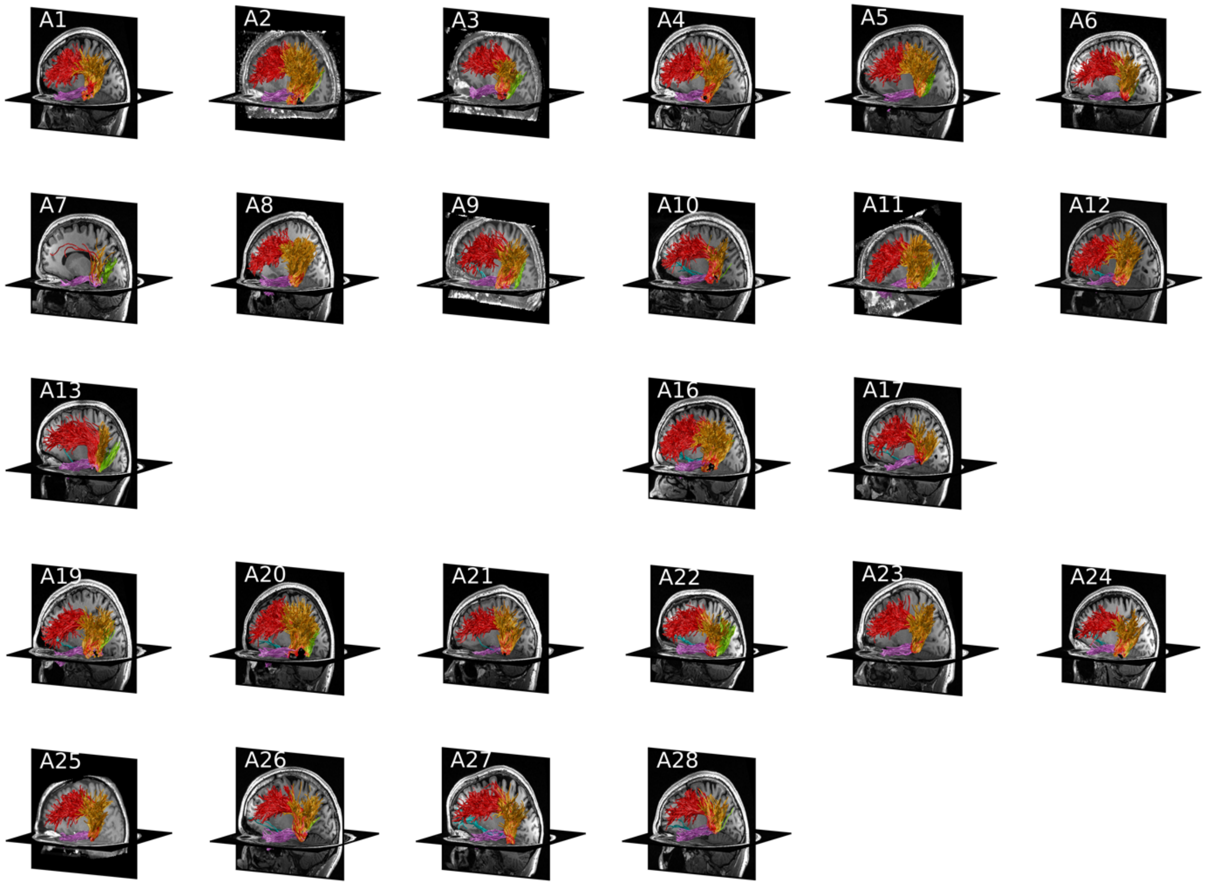


**Supplemental Figure 11. Functionally defined white matter tracts for mOTS-words in individual adult participants (ordered by age).** Upper left of each image: A*:* adult participant, *number:* adult's ranking youngest to oldest. *Black:* functional ROI: mOTS-words. Fascicles: *blue:* inferior frontal occipital fasciculus, *magenta:* inferior longitudinal fasciculus, *red:* arcuate fasciculus, *orange:* posterior arcuate fasciculus, *green:* vertical occipital fasciculus.

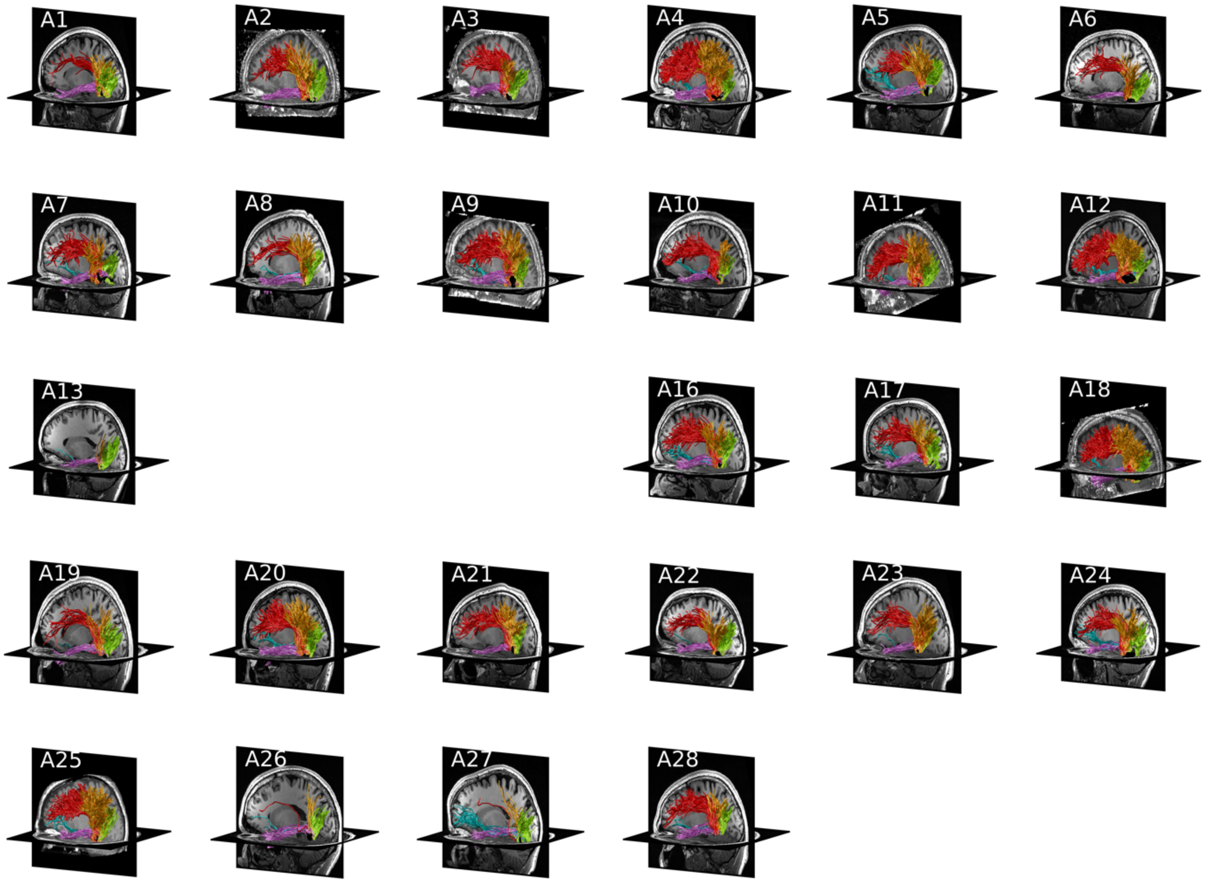


**Supplemental Figure 12. Functionally defined white matter tracts for pOTS-words in individual adult participants (ordered by age).** Upper left of each image: A*:* adult participant, *number:* adult's ranking youngest to oldest. *Black:* functional ROI: pOTS-words. Fascicles: *blue:* inferior frontal occipital fasciculus, *magenta:* inferior longitudinal fasciculus, *red:* arcuate fasciculus, *orange:* posterior arcuate fasciculus, *green:* vertical occipital fasciculus.

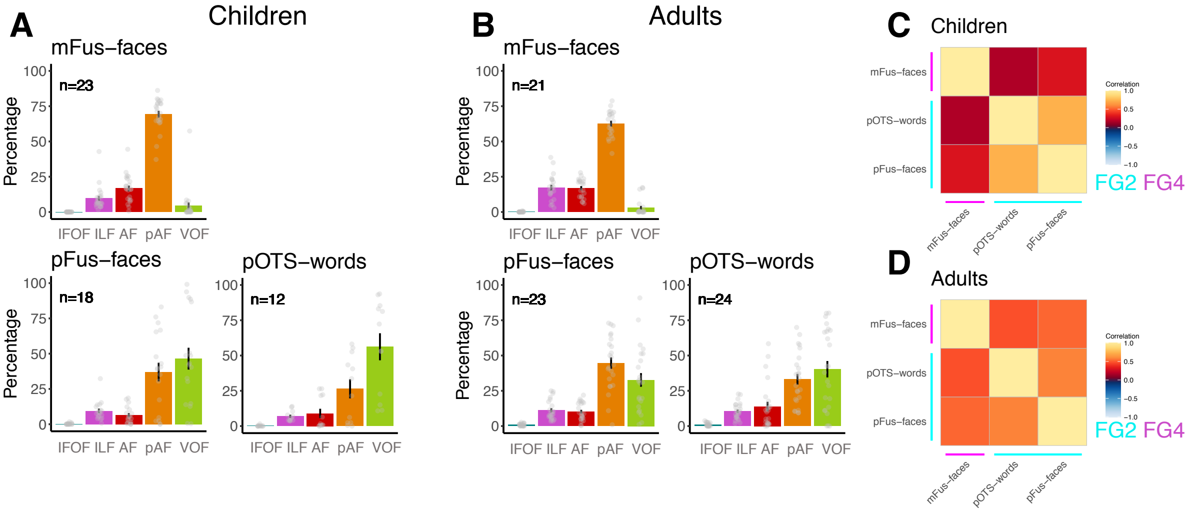


**Supplemental Figure 13. Fascicle connectivity profiles in the right hemisphere.** (A,B) Histograms showing the percentage of connections to major fascicles for each of the functional regions in children (A) and adults (B). *Error bars:* standard error of the mean. *Dots*: individual participant data. *IFOF:* inferior fronto-occipital fasciculus; *ILF:* inferior longitudinal fasciculus; *AF:* arcuate fasciculus; *pAF:* posterior arcuate fasciculus; *VOF:* vertical occipital fasciculus. (C) Correlation matrices depicting the average pairwise within-subject correlation between fascicle connectivity profiles for children. (D) Correlation matrices depicting the average pairwise within-subject correlation between fascicle connectivity profiles for adults. In C,D: Lines depict cytoarchitectonic area (*cyan:* FG2, *magenta:* FG4).

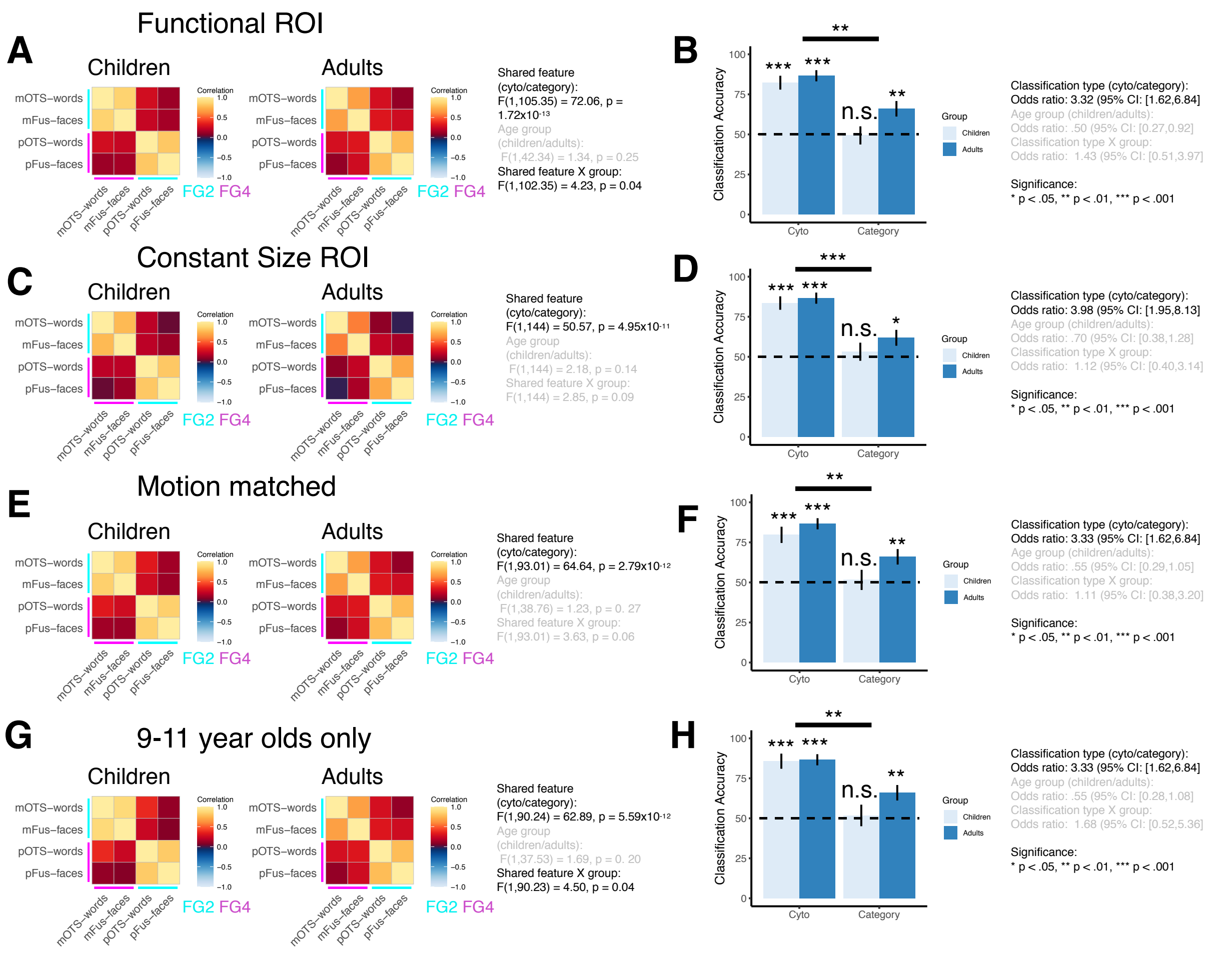


**Supplemental Figure 14. Control analyses for fascicle connectivity profiles.** (A,C,E,G) Correlation matrices depicting the average within-subject pairwise correlation between fascicle connectivity profiles of ventral ROIs in children (left) and adults (right). *Acronyms: mFus-faces:* mid fusiform face-selective region. *pFus-faces:* posterior fusiform face-selective region. *mOTS-words:* mid occipitotemporal sulcus word-selective region. *pOTS-words:* posterior occipitotemporal sulcus word-selective region. (B,D,F,H) Bar graphs showing the average classification accuracy for predicting the cytoarchitecture, category-selectivity, and ROI from endpoint connectivity profiles. *Light blue:* Children, *dark blue:* adults. *Error bars:* standard error of the mean. *Dotted line:* chance level. (A,B) Correlation matrix and classification accuracy for the functional ROIs presented in the main paper. (C,D) Correlation matrix and classification accuracy for 3mm constant disk ROIs centered on each subject's functional ROI. (E,F) Correlation matrix and classification accuracy excluding five child participants such that motion is matched across age groups. (G,H) Correlation matrix and classification accuracy including a subset of child participants in a more restricted age range (9-11 years old).

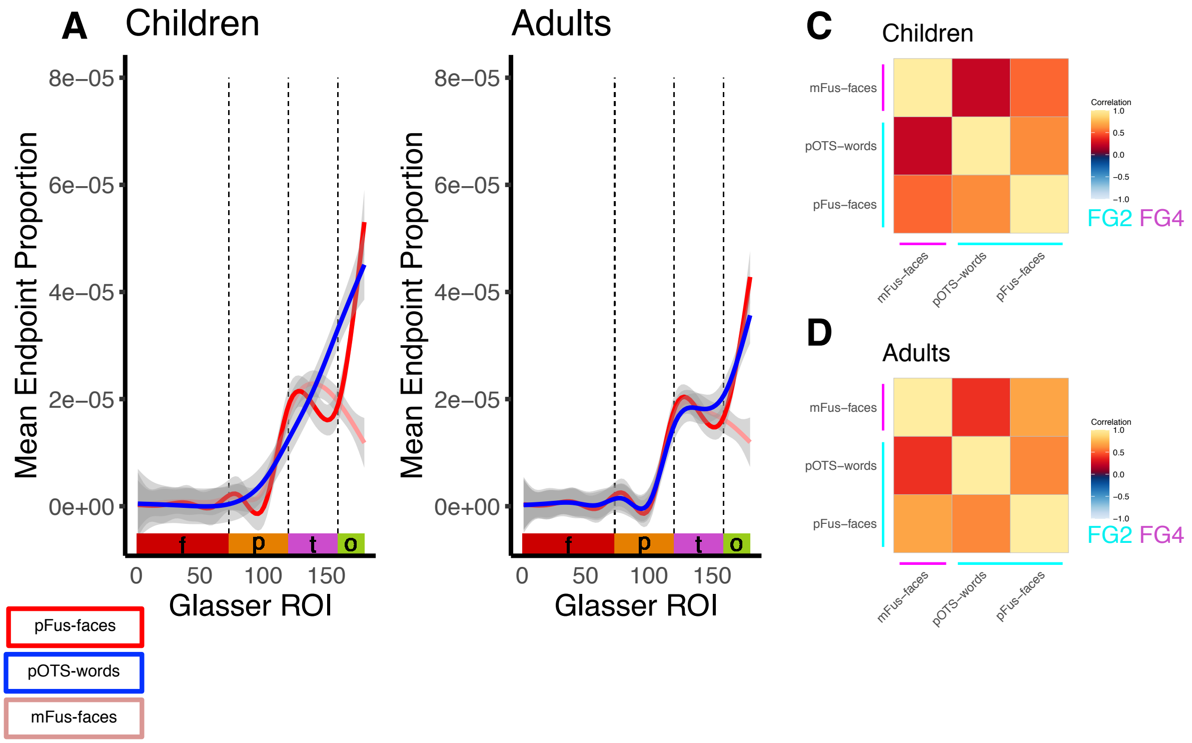


**Supplemental Figure 15. Quantification of endpoint connectivity profile of face- and word-selective regions in children and adults in the right hemisphere**. (A,B) Mean endpoint proportion across 180 Glasser ROIs for the VTC word and face-selective regions in children (A) and adults (B). Line color indicates the functional ROI; *red:* pFus-faces; *blue:* pOTS-words; *light red:* mFus-faces. Shaded area: standard error of the mean. *X-axis:* Glasser Atlas ROI number arranged by lobe: *red:* frontal (f); *orange:* parietal (p); *magenta:* temporal (t); *green:* occipital (o). *Vertical dashed lines:* lobe boundaries. (C,D) Correlation matrices depicting the average within-subject pairwise correlation between endpoint connectivity profiles of ventral face and word-selective ROIs in children (C) and adults (D). *Acronyms: mFus-faces:* mid fusiform face-selective region*. pFus-faces:* posterior fusiform face-selective region. *pOTS-words:* posterior occipitotemporal sulcus word-selective region.

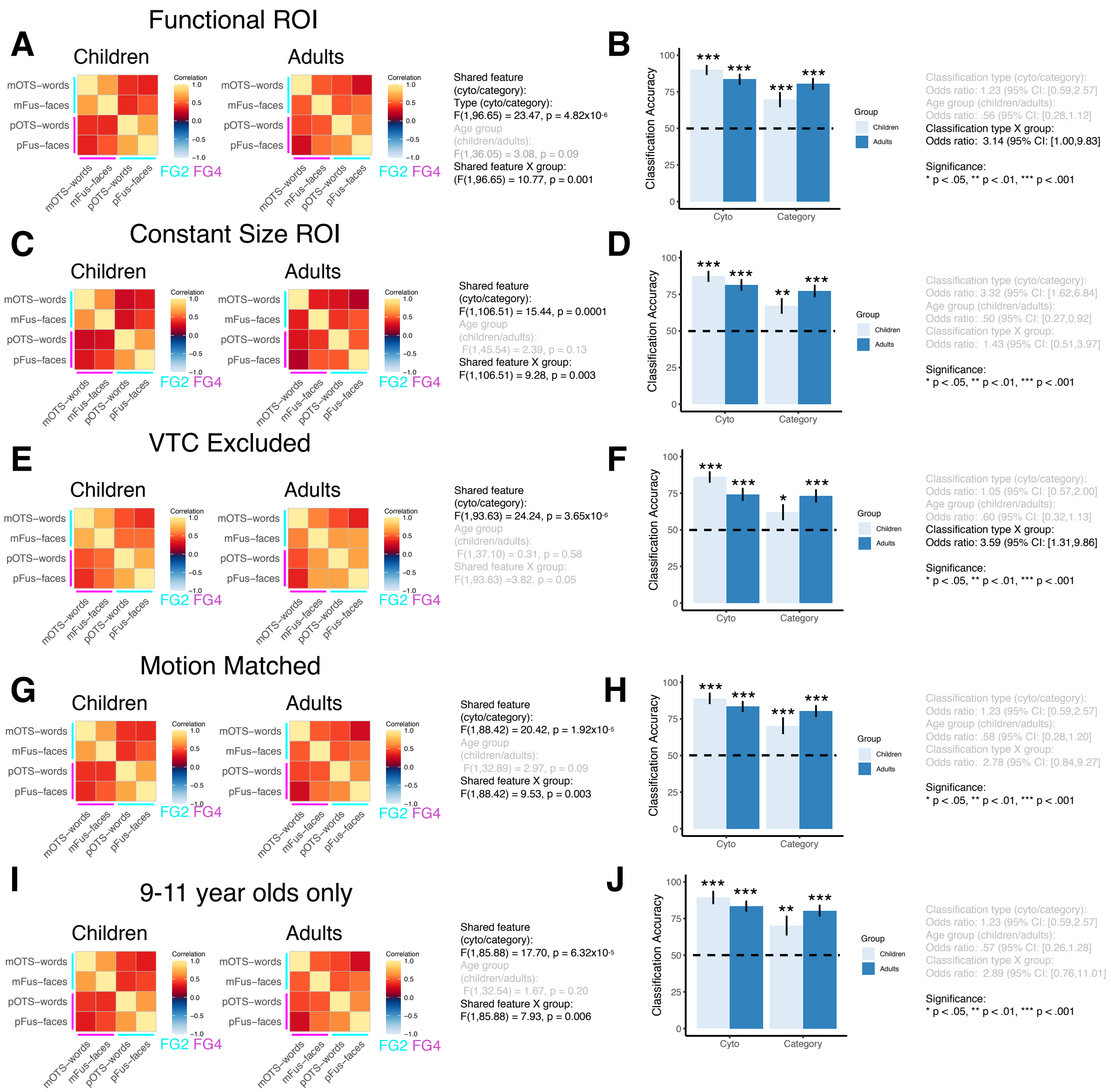


**Supplemental Figure 16. Endpoint connectivity profiles are correlated with connectivity profiles within the same cytoarchitectonic area.** (A,C,E,G,I) Correlation matrices depicting the average within-subject pairwise correlation between endpoint connectivity profiles of ventral ROIs in children (left) and adults (right). *Acronyms: mFus-faces:* mid fusiform face-selective region. *pFus-faces:* posterior fusiform face-selective region. *mOTS-words:* mid occipitotemporal sulcus word-selective region. *pOTS-words:* posterior occipitotemporal sulcus word-selective region. (B,D,F,H,J) Bar graphs showing the average classification accuracy for predicting the cytoarchitecture and category-selectivity from endpoint connectivity profiles. *Light blue:* Children, *dark blue:* adults; *Error bars:* standard error of the mean. *Dotted line:* chance level. (A,B) Correlation matrices and classification accuracy on connectivity profiles of functional ROIs across all 180 Glasser ROIs. (C,D) Correlation matrices and classification accuracy on connectivity profiles of constant size 3mm disk ROIs centered on each participant's functional ROIs across all 180 Glasser ROIs. (E,F) Correlation matrices and classification accuracy on connectivity profiles of functional ROIs calculated over 169 Glasser ROIs excluding 11 VTC ROIs. (G,H) Correlation matrices and classification accuracy on connectivity profiles of functional ROIs excluding five child participants such that motion is matched between the age groups. (I,J) Correlation matrices and classification accuracy on connectivity profiles of functional ROIs including a subset of child participants within a tighter age range (9-11 years old).

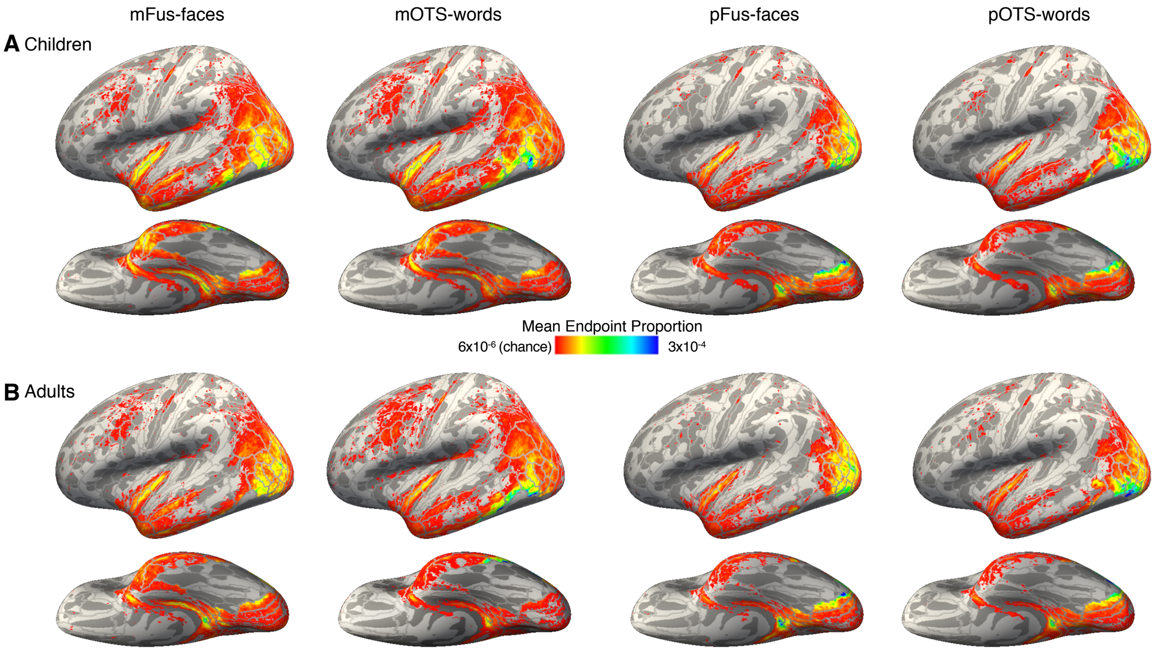


**Supplemental Figure 17. Endpoint connectivity profile of face- and word-selective with VTC excluded.** Average endpoint connectivity profile of mFus-face, pFus-faces, mOTS-words, pOTS-words in children (A) and adults (B). Data are shown on the fsaverage inflated cortical surface. *Color map:* endpoint proportion at each vertex. *Threshold value of maps:* chance level if all endpoints were evenly distributed across the brain. *Gray outlines:* Glasser ROIs.

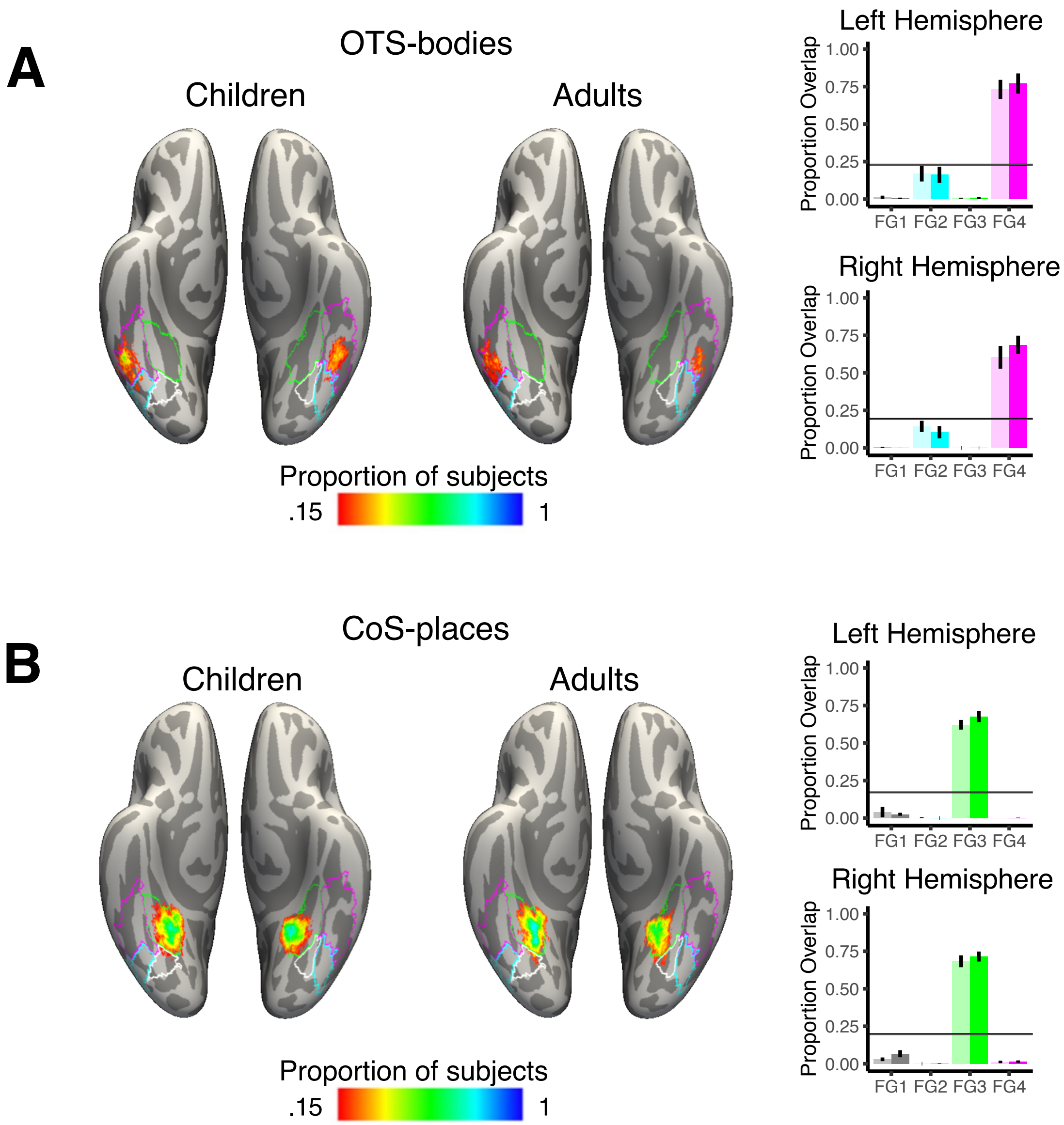


**Supplemental Figure 18. Overlap between functional ROIs and cytoarchitectonic ROIs for OTS-bodies and CoS-places.** Each brain depicts a probabilistic map of a functional ROI in relation to the boundaries of the maximal probability map (MPM of each cytoarchitectonic area on the fsaverage cortical surface. *Colormap*: the proportion of subjects at each vertex for which this vertex is within the functional ROIs. Outlines: boundaries of the cytoarchitectonic ROIs (FG1: white, FG2: cyan, FG3: green, FG4: purple). Children's data are shown on the left and adults' are on the right. Bar graphs depict the proportion of overlap between each functional ROI and each cytoarchitectonic area. *Error bars:* standard error across subjects. *Lighter colors:* children, *darker colors:* adults. *Upper:* left hemisphere, *lower:* right hemisphere.
